# Memory Regulatory T Cells Reprogram into Protective Tfh-like Effectors in Recurrent Malaria

**DOI:** 10.1101/2025.10.15.682462

**Authors:** Nana Appiah Essel Charles-Chess, Anthony A. Ruberto, Carson Bowers, Nyamekye Obeng-Adjei, Matthew Richard Hansen, Disha Bangalore Renuka Prasad, Boubacar Traore, Kimberly D. Klonowski, Peter D. Crompton, Samarchith P. Kurup

**Affiliations:** Department of Cellular Biology, University of Georgia, Athens, GA, USA; Center for Tropical and Emerging Global Diseases, University of Georgia, Athens, GA; Institute of Bioinformatics, University of Georgia, Athens, GA, USA; Malaria Infection Biology and Immunity Section, Laboratory of Immunogenetics, Division of Intramural Research, National Institute of Allergy and Infectious Diseases, US National Institutes of Health, Rockville, Maryland, USA; Mali International Centre of Excellence in Research, University of Sciences, Techniques and Technologies of Bamako, Bamako, Mali

## Abstract

Although people living in malaria-endemic areas experience repeated infections with *Plasmodium*, the role of regulatory T cells (Tregs) in recurrent malaria remains poorly understood. During a primary infection with *Plasmodium*, Tregs suppress protective immunity by inhibiting germinal center (GC) reactions, thereby impeding the control of parasitemia. In contrast, we demonstrate here that memory Tregs (mTregs) remaining after the clearance of initial *Plasmodium* infection acquire protective functions upon recall. Relying on longitudinal studies in humans and mice, we show that mTregs undergo antigen-driven expansion and inflammation-induced epigenetic reprogramming during reinfection to transition from Foxp3^+^ immunosuppressive cells to Bcl6^+^ follicular T helper (Tfh)-like effectors. These mTreg-derived Tfh-like cells enhance GC responses and the generation of *Plasmodium*-specific antibodies, ultimately facilitating *Plasmodium* control. Precluding such mTreg-to-Tfh differentiation abolished protection. Our findings reveal a previously unrecognized adaptive plasticity in canonical mTregs that enables a context-dependent functional switch from immunoregulatory to protective effectors during recurrent infections.

## Introduction

A central tenet of adaptive immunity is the expansion in numbers of lymphocytes upon antigen encounter, followed by contraction into a memory pool that enables rapid recall responses upon re-exposure to the same antigen^1^. A key aspect of these responses is immunoregulation by regulatory T cells (Tregs), which are a specialized subset of helper T (Th) cells that prevent excessive immune activation and immunopathology^1^. Tregs specific to self or non-self-antigens can persist as memory Tregs (mTregs) beyond initial immune responses and can undergo recall activation, proliferation, and differentiation into potent suppressors, dampening proinflammatory responses^2^. Such cells are considered essential for mitigating systemic and organ-specific immunopathology when repeated antigen exposures occur, such as in the cases of seasonal flu, recurring skin infections, allergies, and repeat pregnancies^2, 3, 4, 5, 6, 7^.

Malaria, caused by *Plasmodium* parasites, presents a similar scenario of repeated antigen exposures. Over 250 million new clinical cases of malaria are reported each year, largely from malaria-endemic areas, with repeat infections accounting for over 90% of the cases^8, 9, 10, 11, 12^. CD4 T cells are essential for protection from acute malaria, with conventional Th1 cells mediating cytokine-driven parasite clearance and T follicular helper (Tfh) cells supporting the generation of protective antibody responses^13^. Studies in humans naturally or experimentally infected with *Plasmodium* have consistently shown that Tregs also expand during the acute phase of malaria, with higher Treg frequencies correlating with worse disease outcomes^14, 15, 16^. Functional or numerical depletion of Tregs facilitates better control of primary *Plasmodium* infection in mice^13, 17^. This has led to the notion that Tregs impede immunity to malaria. In contrast, the role of CD4 T cells in recurrent malaria remains poorly understood. Recent studies, including in malaria, have revealed diverse recall dynamics among distinct memory CD4 T cell subsets^18, 19, 20, 21^. For example, clonally restricted type I regulatory (T_R_1) T cells exhibited tangible recall expansion upon antigen re-encounter in humans^21^. In mice, while the Th1 cells proliferated rapidly, germinal center (GC) Tfh cells remained refractory to reactivation during re-infection^18^. However, the functional fidelity of any such recalled CD4 T cell subsets remains largely unknown.

There is emerging evidence for distinct CD4 T cell subsets exhibiting functional plasticity in diverse inflammatory environments^22^. Epigenetic profiling of Tfh cells has suggested potential plasticity with various other Th subsets. For example, late-stage GC Tfh cells were recently shown to upregulate Foxp3 and other Treg-associated functional markers^23, 24^. Tregs within the skin, Peyer’s patches, or atherosclerotic vessels have also been observed to progressively acquire a Tfh-like phenotype^25, 26, 27, 28, 29^. In the context of *Plasmodium* reinfection, memory Th1 cells can adopt an immunosuppressive phenotype, marked by IL-10 production and the upregulation of inhibitory molecules^18^. This raises the question whether such plasticity extends to mTregs as well, particularly in the context of repeat infections. This is a crucial knowledge gap, considering that long-lived Tregs are increasingly being recognized as potential therapeutic targets in recurring or persistent diseases such as flu, malaria, autoimmune diseases, and cancer^30, 31, 32, 33, 34^. If naïve and memory Tregs serve distinct functions, immunotherapies seeking to modify Treg responses would have to be tailored to the specific context to produce positive outcomes. Recurrent malaria provides a unique opportunity to interrogate how repeated waves of antigen exposures and intense inflammation reshape Treg memory, both functionally and conceptually.

Here, we investigate the generation, maintenance, and recall of mTregs in recurrent infections. Relying on longitudinal studies in humans and mice repeatedly exposed to *Plasmodium*, *Listeria,* or flu virus, we show that canonical mTregs downregulate Foxp3 and transition into Tfh-like effectors during reinfection, promoting antibody-mediated protection. This switch is associated with broad, inflammation-mediated epigenetic reprogramming, including in their lineage-defining loci. Blocking this transition prevented mTregs from conferring protection against recurrent malaria. Our findings redefine mTregs as a population of dynamically plastic cells, challenging the current paradigm of their stable functional identity through recall.

## Results

### Establishment of Treg memory following *Plasmodium* infection

To investigate whether *Plasmodium* infection induces lasting regulatory T cell memory, we analyzed peripheral blood in a cohort of Malian children at least six months after the resolution of *P. falciparum* malaria following anti-malarial drug treatment, and with no other reported febrile illnesses in the interim (**Figure 1A, S1A**)^11^. These children maintained significantly higher frequencies of memory Tregs (mTregs; Helios^+^ CTLA-4^+^ FOXP3^+^ CD4 T cells)^35, 36^ compared to the malaria-naïve U.S controls (**Figure 1B**), suggesting that *Plasmodium* infection induces persistence of mTregs in humans.

**Figure 1.**
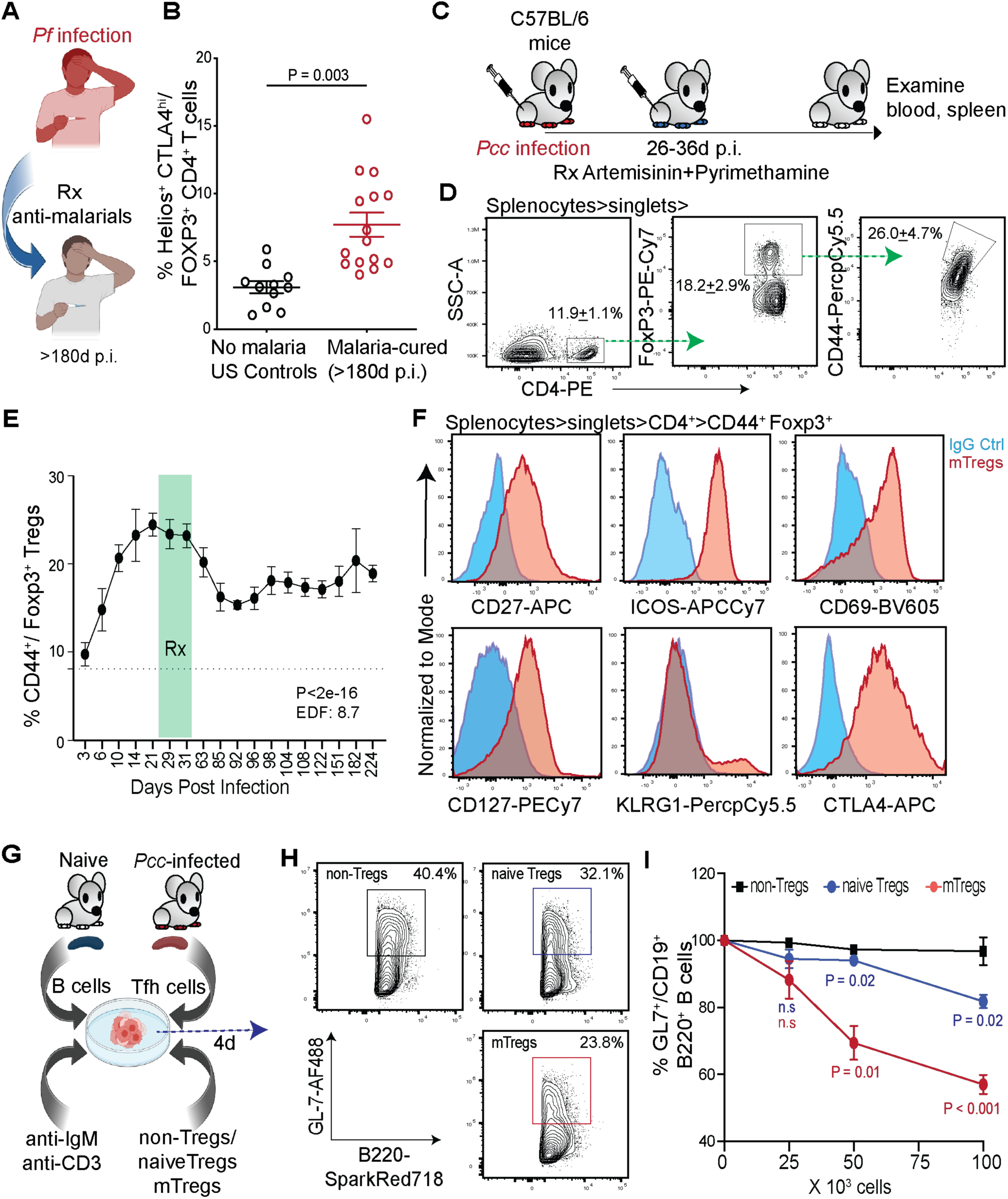
Long-term persistence of memory Tregs following *Plasmodium* infection in humans and mice. (A-B) Malian children diagnosed with naturally acquired malaria were treated with anti-malarial drugs to achieve parasitological cure and their peripheral blood was examined ≥180d post-infection (p.i.) (A). Relative frequencies of Helios^+^ CTLA4^hi^ FOXP3^+^ CD4^+^ CD3^+^ mTregs in the peripheral blood of malaria-naïve US adults or malaria-experienced Malian children. Each dot represents an individual, data analyzed using student t-test, yielding the indicated *P*-value (B). (C-D) Experimental scheme: B6 mice infected with *Plasmodium chabaudi chabaudi* (*Pcc*) were treated with Artemisinin-Pyrimethamine combination and the spleens examined ≥90d p.i (C). Representative flow-plots depicting splenic Foxp3^+^ CD44^+^ CD4 T cell (mTreg) frequencies and the gating strategy (D). Numbers inset indicate frequencies of the gated populations presented as mean ± standard deviation (SD) from 1 of 3 replicate experiments, n=3. (E) Temporal kinetics of splenic mTreg frequencies in *Pcc*-exposed B6 mice generated as in (C). Vertical green bar indicates the anti-malarial treatment window, combined data presented as mean ± standard error of mean (s.e.m.) from ≥3 separate experiments initiated with ≥3 mice/group. Statistical modeling of temporal trends in mTreg frequencies was performed using a generalized additive model (GAM) with a smooth term for time (d p.i.) to identify the effect of time on mTreg frequencies, yielding the indicated *P*-value and estimated degrees of freedom (EDF) across the 0-224d window. (F) Representative histograms (red) depicting expression of the indicated markers in mTregs identified as in (D); blue histograms show IgG staining controls. (G-I) Experimental scheme: *In vitro* germinal center (GC) reaction composing B cells and follicular T helper (Tfh) cells obtained from naïve and *Pcc*-exposed B6 mice (28d p.i.), respectively, were seeded with increasing numbers of splenic naïve Tregs (Foxp3^+^ CD4^+^, from naïve B6 mice), or non-Tregs (Foxp3^-^ CD44^+^ CD4^+^) or mTregs (Foxp3^+^ CD44^+^ CD4^+^) obtained from *Pcc*-exposed B6 mice (181d p.i.) in the presence of anti-IgM and anti-CD3 antibodies (G). Representative flow plots showing GL7^+^ CD19^+^ B220^+^ GC B cell frequencies at 4d of co-incubation with numbers inset indicating gated cell frequencies (H). Cumulative data normalized for input shown in line graphs (I), presented as mean ± SD from 1 of 2 separate experiments with 3 technical replicates/experiment, analyzed using 2-way ANOVA with Dunnett’s correction comparing the color-coded groups (blue and red) with the non-Treg group, yielding the indicated *P*-values (i). n.s = *P*>0.05.

To reproduce the infection-cure scenario described above in mice, and to gain mechanistic insights into mTreg cell generation and function, we employed the murine *P. chabaudi chabaudi* (*Pcc*)-infection model. *Pcc* broadly recapitulates human *P. falciparum* infection biology in mice and can establish repeat infections, as in humans^37^. Drug-cured *Pcc*-experienced mice retained phenotypic mTregs (defined as CD44^+^ Foxp3^+^ CD4 T cells)^2^ in circulation and lymphoid tissues for over six months post-infection (p.i.), as observed in the Malian cohort (**Figure 1C-E**). Such mTregs expressed canonical markers of memory and phenotypically resembled the mTregs retained in *Listeria monocytogenes* (*Lm*) or influenza A virus (IAV)-exposed mice (**Figures 1F, S1B, S1C**)^7, 38, 39^.

The mTregs in *Pcc*-experienced mice (180d p.i.) also expressed CD25 and Neuropilin-1 (Nrp-1) at levels similar to that of naïve Tregs (**Figure S1D**), suggesting their preserved regulatory capacity^40^. These mTregs also expressed T-bet, a transcription factor associated with long-term survival and immunosuppressive capacity in Tregs, mirroring the T-bet^+^ signature of the Tregs in malaria-exposed Gambian individuals, at the end of malaria transmission season **(Figure S1E)**^2, 41^. Functional assays further confirmed the suppressive nature of such mTregs in mice. In a standard *in vitro* immunosuppression assay, sorted mTregs derived from *Pcc*-experienced FOXP3-DTR^GFP^ mice suppressed the proliferation of chicken ovalbumin antigen (OVA)-specific (OT-I) T cells co-incubated with OVA-pulsed dendritic cells (DCs) in a dose-dependent manner (**Figure S1F-G**). The mTregs derived from *Pcc*-experienced mice also significantly inhibited the generation of GC B cell responses *in vitro* (**Figure 1G-I**). Together, these findings demonstrated that *Plasmodium* infection induces a population of phenotypically consistent and functionally competent mTregs that persist long-term in both humans and mice.

### Memory Tregs associated with protection from *Plasmodium* reinfection in humans and mice

Based on how T cells are expected to retain their original functions during recall, we predicted that mTregs, like the effector Tregs in a primary infection, would impede protection during *Plasmodium* reinfection. However, in longitudinal studies conducted in humans, we discovered that higher pre-season mTreg frequencies correlated with reduced *P. falciparum* parasitemia following natural infection in the subsequent malaria transmission season (**Figure 2A-B**). This protective association also correlated with age and the associated increase in malaria exposures^11, 33^, suggesting that recurring infections might drive mTreg accumulation and functional adaptations following repeated antigen-driven recalls in humans.

**Figure 2.**
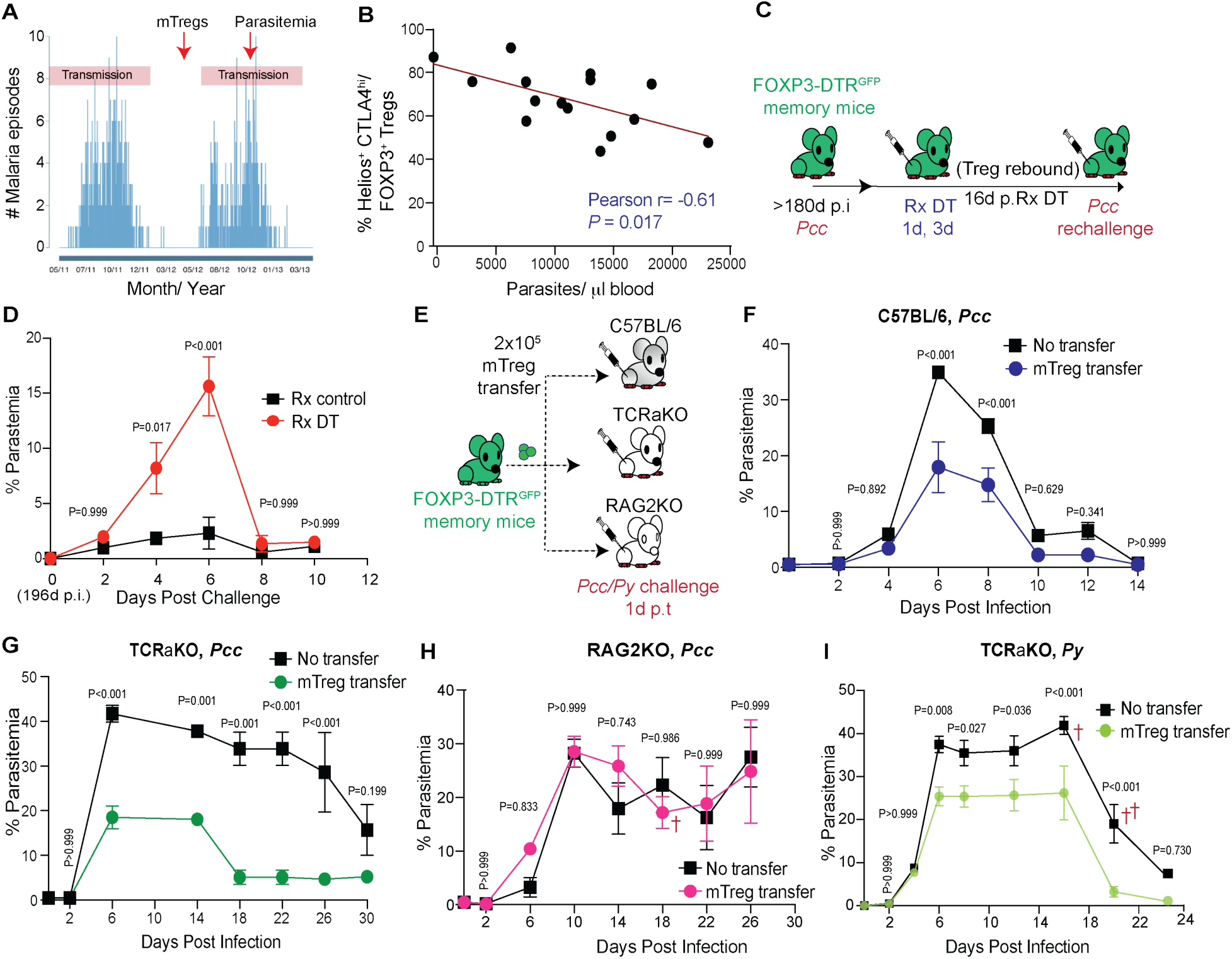
Memory Tregs associate with decreased *P. falciparum* parasitemia in humans and confer protection against *Plasmodium* challenge in mice. (A-B) Histogram representing the seasonal incidence of clinical malaria at the study site and the timing of collection of peripheral blood to quantify mTregs and parasitemia in the Malian cohort (A). Scatter plot showing correlation between pre-existing mTreg frequencies before the malaria season and parasitemia during the first febrile malaria episode of the ensuing malaria-transmission season. Each dot represents an individual, Pearson r and *P*-values obtained through regression analysis (B). (C-D) Experimental scheme: *Pcc*-experienced FOXP3-DTR^GFP^ mice (180d p.i.) treated with diphtheria toxin (DT, i.p., 181d and 184d) were rechallenged with *Pcc* (2x10^6^ iRBCs/mouse) at 196d p.i. (C). Temporal kinetics of parasitemia in the DT- or control (PBS)-treated mice (D). (E-I) Experimental scheme: 2x10^5^ mTregs flow-sorted from *Pcc*-experienced FOXP3-DTR^GFP^ mice (180d p.i.) were adoptively transferred to B6, TCRaKO, or RAG2KO recipient mice, which were then challenged with *Pcc* or *Py* (5x10^5^ iRBCs/mouse) at 1d post-transfer (p.t.) (E). Temporal kinetics of parasitemia in the B6 (F), TCRaKO (G, I), or RAG2KO (H) recipients. Representative data from 1 of 3 experiments presented as mean ± SD, analyzed using 2-way ANOVA with Sidak correction for multiple comparison, yielding the indicated *P*-values (D, F-I). Experiments initiated with 3 (D, F-H) or 5 (I) mice; ^†^Death of 1 mouse/group.

To mechanistically interrogate the protective role of these mTregs, we leveraged the FOXP3-DTR^GFP^ mouse model, in which treatment with diphtheria toxin (DT) ablates all Foxp3-expressing cells, allowing precise temporal control of Treg populations^42^. To isolate the role of mTregs, we treated *Pcc*-experienced memory FOXP3-DTR^GFP^ mice with DT to deplete all Tregs, including the mTregs, before permitting the replenishment of naïve Tregs to the pre-depletion levels (**Figure 2C, Figure S2A**). Rechallenge of these mice resulted in significantly higher *Pcc* parasitemia than the mTreg-sufficient control mice (**Figure 2D**). Of note, immunocompetent mice are expected to fully control the *Pcc* infection eventually^37^.

TCRaKO mice, which lack T cells, serve as a more stringent model to determine the protective roles of T cells against malaria^43^. It is well-established that adoptive transfer of the memory helper T (mTh) cell subset is sufficient to confer immunity to *Plasmodium* infection in TCRaKO mice^19^. However, selective depletion of mTregs from the transferred mTh compartment compromised protection in the *Pcc*-infected TCRaKO recipients (**Figure S2B-C**). Remarkably, the transfer of mTregs isolated from *Pcc*-experienced mice alone was sufficient to offer protection from *Pcc* challenge in both wild-type and TCRaKO recipients (**Figure 2E-G**). Together, these data suggested that mTregs are both necessary and sufficient to protect mice from *Pcc* infection. But the transfer of mTregs failed to protect RAG2KO mice (**Figure 2H**). Transfers of even up to 3x the effective dose of mTregs in TCRaKO mice failed to protect RAG2KO mice from *Pcc* infection (data not shown), implying that B cells are required for the protection conferred by mTregs.

This mTreg-mediated protection also showed unexpected breadth and specificity. The mTregs obtained from IAV-experienced, but not *Pcc*-experienced mice conferred protection against IAV challenge (**Figure S3**), consistent with the lack of shared epitopes between IAV and *Plasmodium*. However, *Pcc*-primed mTregs offered cross-protection against the antigenically related, but distinct *Plasmodium* species, *P. yoelii* (**Figure 2I**). This indicated that mTreg-mediated protection is possibly antigen-restricted. Importantly, the adoptive transfer of an equal number of naïve Tregs failed to provide any protection from *Pcc* infection in B6 or TCRaKO mice (**Figure S4**), implying that memory recall and antigen specificity may be critical features of mTreg-mediated protection.

Given the overall unexpected nature of the above findings, we considered whether the protection observed with mTreg transfers may be due to inadvertent co-transfer of other protective T cell subsets such as Th1, Tfh cells, etc. Although we flow-sorted the mTregs at high-fidelity (purity mode), yielding >99% GFP^+^ CD44+ CD4 T cells, there is a chance that flow sorting itself or the adoptive transfer process might alter the mTreg phenotype or function (**Figure S5A**). To rule this possibility out, we depleted the Foxp3-expressing cells from the adoptively transferred FOXP3-DTR^GFP^ mTreg population by treating the recipients with DT before *Pcc* challenge (**Figure S5B**). This reversed the control of *Pcc* in the TCRaKO recipients (**Figure S5C**). As a more stringent test, we employed IFNARKO mice, which are lethally susceptible to *Pcc* infection, as mTreg recipients above. The depletion of Foxp3^+^ cells from the flow-sorted mTregs made the IFNARKO mice just as susceptible to *Pcc* infection, as the no-mTreg-transfer group (**Figure S5D**). Together, these findings reveal a fundamental shift in Treg function during recurrent malaria-from immunosuppression in primary infection to a B cell-dependent, potentially antigen-restricted protection, during reinfection.

### Memory Tregs augment *Plasmodium*-specific humoral immunity through germinal center expansion

To investigate the mechanism of mTreg-mediated protection, we transferred congenically distinct mTregs from *Pcc*-experienced donor mice into naïve B6 recipients, followed by challenge with *Pcc* (**Figure 3A**). Remarkably, the recipients exhibited broad enhancements in adaptive immune responses with significantly increased numbers of splenic activated CD4 T cells, Tfh cells, B cells, GC B cells, and GC plasmablasts, when compared to the control mice that did not receive mTregs (**Figure 3B-G**). Correspondingly, the mTreg recipients also generated larger splenic GCs covering greater overall splenic volumes (**Figure 3H-K**). These cellular responses correlated with elevated *Plasmodium*-specific IgM and IgG titers (**Figure 3L-M**) in the recipient mice, directly linking the mTreg activity to humoral immunity. These findings confirmed that mTregs amplify antigen-specific immunity to *Plasmodium*.

**Figure 3.**
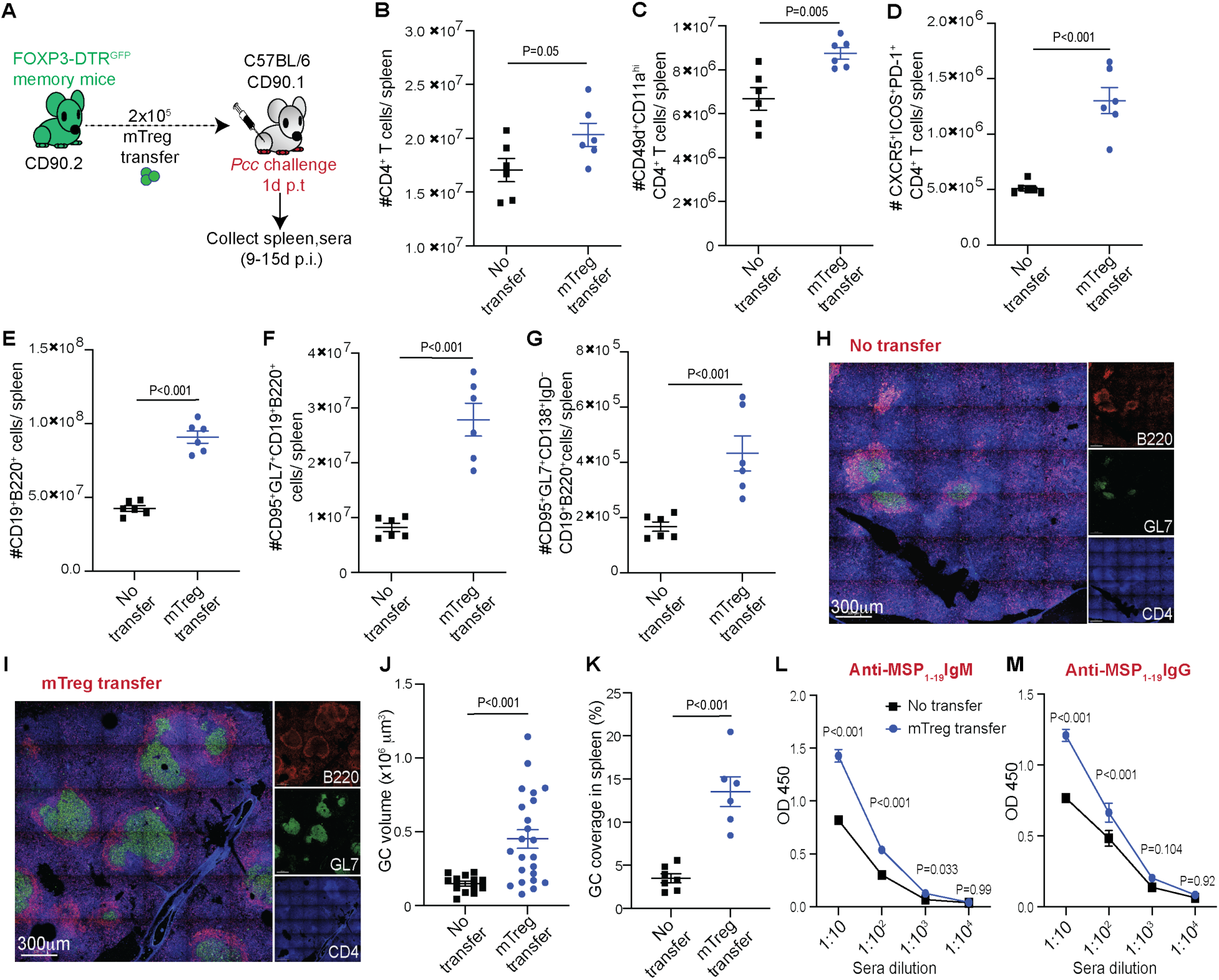
Memory Tregs promote stronger T helper and B cell responses in *Plasmodium*-infected mice. (A) 2x10^5^ mTregs derived from *Pcc*-experienced FOXP3-DTR^GFP^ mice (≥180d p.i.) were adoptively transferred to congenically distinct B6 recipients, which were then challenged with *Pcc* (5x10^5^ iRBCs/mouse). Media inoculated as ‘no transfer’ controls. (B-G) Scatter plots indicating total numbers of splenic CD4 T cells (B), *Plasmodium*-specific CD4 T cells (C), Tfh cells (D), B cells (E), GC B cells (F), or GC plasmablasts (G) in the recipients, 15d p.i. Combined data from 2 experiments presented as mean ± s.e.m.; each dot represents a recipient mouse. (H-I) Representative pseudocolored confocal images showing GC reactions in recipient spleens without (H) or with (I) mTreg transfer, 15d p.i. Scatter plots summarize total GC volume (J) or GC coverage (K) in the spleen sections. Combined data presented as mean ± s.e.m. with each dot representing a GC (J) or spleen section (K) from 3 experiments, n=3 mice. (L-M) Relative levels of *Plasmodium* MSP_1-19_–specific IgM (L) or IgG (M) antibody titers in the sera of the recipients (9d p.i.) with or without mTreg transfer. Data in B-G, J-K analyzed using t-tests, L-M analyzed using 2-way ANOVA with Sidak correction, yielding the indicated *P*-values.

### Recall of memory Tregs associated with inflammation-driven loss of Foxp3 expression

To assess the proliferative responses in mTregs, recipients of *Pcc*-experienced mTregs were challenged with *Pcc*, while under BrdU treatment. A substantial proportion of the transferred mTregs incorporated BrdU, revealing antigen-driven recall expansion (**Figure 4A-B**). Unexpectedly though, the vast majority of the transferred cells also appeared to have simultaneously lost their Foxp3 expression (**Figure 4C)**. These Foxp3-negative descendants of mTregs, which we term ‘xTregs’ (short for ex-mTregs), were detectable only following *Pcc* challenge, and remained undetectable in the uninfected recipients (data not shown). Also, the transition of mTregs to Foxp3^-^ xTregs was not unique to *Plasmodium* infection. The mTregs generated in *Lm* or IAV infected mice downregulated their Foxp3 expression following re-exposure to *Lm* or IAV respectively, in the recipient mice (**Figure S6**).

**Figure 4.**
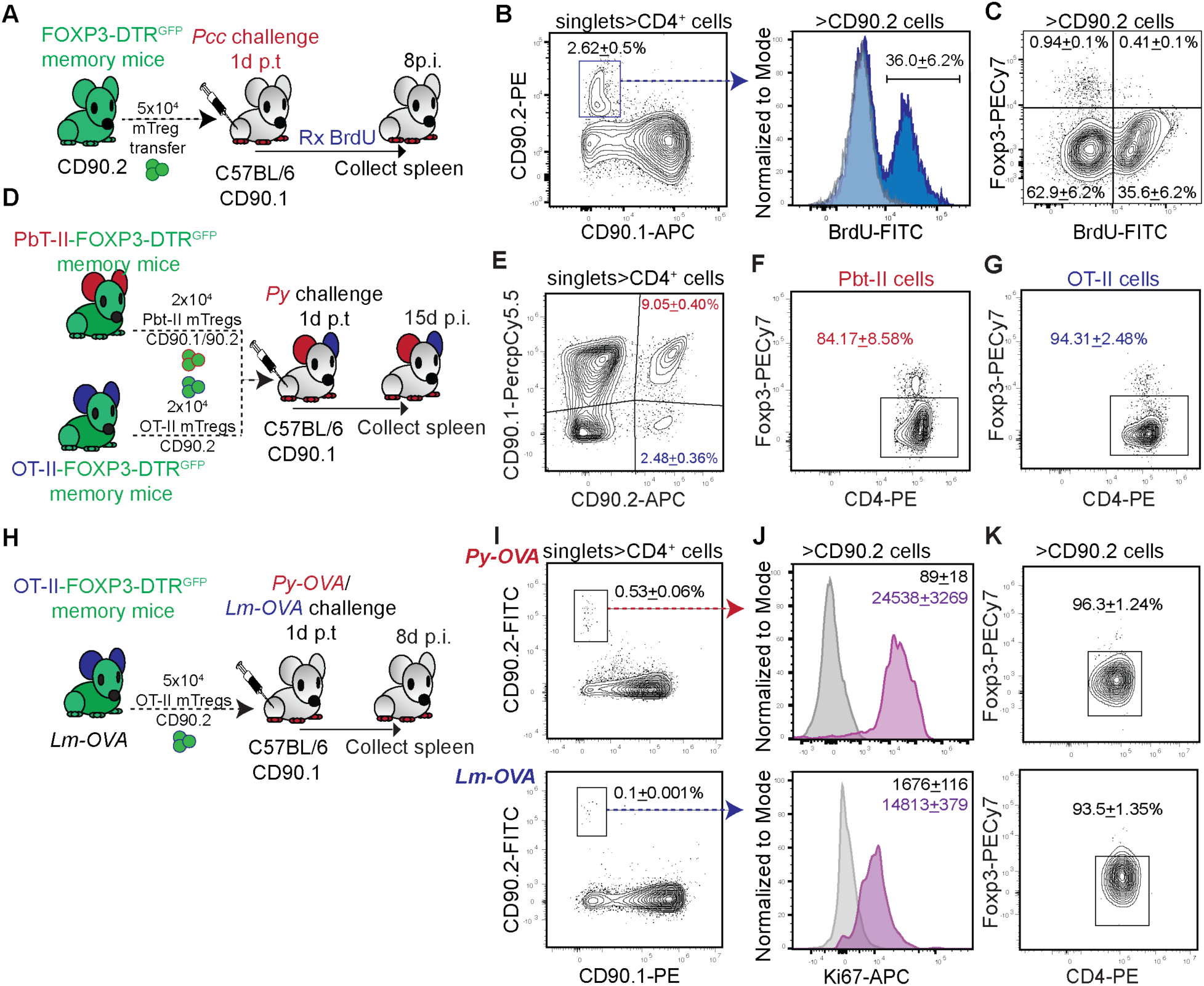
Memory Treg recall associated with downregulation of Foxp3 expression. (A-C) Experimental scheme: 5x10^4^ mTregs derived from *Pcc*-experienced FOXP3-DTR^GFP^ mice (CD90.2, ≥180d p.i.) were adoptively transferred to congenically distinct (CD90.1) B6 mice treated with BrdU and challenged with *Pcc* (5x10^5^ iRBCs/mouse) (A). Representative flow plots and histograms indicating BrdU incorporation (B) and Foxp3 expression (C) in the donor cells recovered from recipient spleen, 8d p.i. Numbers inset show the gated cell frequencies presented as mean ± SD from 1 of 3 identical experiments, n=3 recipients. (D-G) 2x10^4^ each of PbT-II (CD90.1/90.2) or OT-II (CD90.2) mTregs (98d p.i.) generated in *Py- OVA*-exposed PbT-II-FOXP3-DTR^GFP^ or OT-II-FOXP3-DTR^GFP^ mice, respectively, were adoptively co-transferred to congenically distinct B6 mice (CD90.1), which were then challenged with *Py* (2.5x10^5^ iRBCs/mouse), and analyzed at 15d p.i. (D). Representative flow plots depicting relative frequencies of PbT-II or OT-II cells (E) or their Foxp3 expression (F, G) in the recipients. Numbers inset indicate the frequencies of the gated populations presented as mean ± SD from 1 of 2 replicate experiments; n=3 recipients. (H-K) 5x10^5^ OT-II (CD90.2) splenic mTregs (≥180d p.i.) generated from *Lm-OVA*-experienced OT-II-FOXP3-DTR^GFP^ mice were adoptively transferred to congenically distinct B6 mice (CD90.1), which were then challenged with *Pb-OVA* iRBCs (5x10^6^ iRBCs/mouse) or *Lm-OVA* (ActA^-^, 1x10^7^) and analyzed 15d p.i. (H). Representative flow plots or histograms depicting relative frequencies (I) of splenic OT-II cells and their expression of Ki67 (J) or Foxp3 (K) following *Py-OVA* (upper panel) or *Lm-OVA* (lower panel) challenge of the recipients. Numbers inset indicate the frequencies of the gated populations (I, K) or geometric mean fluorescent intensities (gMFI, J) of Ki67 (purple histogram) or IgG ctrl (gray histogram), presented as mean ± SD from 1 of 3 replicate experiments; n=3 recipients. All cells isolated or analyzed from spleens.

To determine the extent to which FOXP3 was downregulated in humans during antigen-specific mTreg recall, we interrogated the existing publicly available single-cell datasets of CD4 T cells obtained from individuals naturally re-exposed to *P. falciparum*^21^. Indeed, several Treg clones expanded in a TCR-restricted manner following *P. falciparum* re-exposure and significantly downregulated their *FOXP3* transcripts (**Figure S7**).

To delineate the roles of cognate antigen and inflammatory signals in driving mTreg-to-xTreg transition in antigenically restricted populations, we generated PbT-II-FOXP3^GFP^ reporter mice by crossbreeding PbT-II TCR-transgenic (containing CD4 T cells specific for *Plasmodium* heat shock protein (Hsp) 90 antigen)^44^ and FOXP3-DTR^GFP^ mice. The naïve PbT-II precursors in PbT-II-FOXP3^GFP^ mice represented 20-50% of the CD4 T cell compartment and following *Py-OVA* (*P. yoelii* expressing OVA) infection, generated an antigen-specific mTreg population comprising ∼10% of the CD4 memory pool at 180d p.i, which phenotypically resembled the polyclonal mTregs from B6 mice (**Figure S8A).** These PbT-II mTregs underwent robust recall proliferation following *Pcc* re-encounter, as indicated by their enhanced Ki67 expression compared to the naïve PbT-II Tregs (**Figure S8B-D**)^45^. Also, as with the polyclonal naïve Tregs, PbT-II naïve Tregs did not downregulate Foxp3 expression following *Pcc* encounter (**Figure S8E-I**).

In similar lines, we developed the OT-II-FOXP3^GFP^ reporter line, by crossbreeding OT-II TCR-transgenic (containing OVA-specific CD4 T cells)^46^ and FOXP3-DTR^GFP^ mice. The OT-II precursors also constituted 20-50% of the CD4 T cell compartment, and following *Py-OVA* infection, generated OT-II mTreg frequencies comprising ∼10% of the CD4 memory pool at 180d p.i, similar to the PbT-II mTregs in PbT-II-FOXP3^GFP^ memory mice (data not shown). Co-transfer of equal numbers of flow-sorted PbT-II and OT-II mTregs to *P. yoelii*-infected recipients resulted in robust PbT-II mTregs expansion through antigen-driven recall, and OT-II mTregs proliferation, possibly through bystander activation (**Figure 4D-E**). However, both PbT-II and OT-II mTregs similarly downregulated their Foxp3 expression in the recipients (**Figure 4F-G**). We confirmed the identities of OT-II (Vα2/Vβ5) and PbT-II (Vα2/Vβ12) cells in the recipients by staining for their respective TCR Vα/Vβ clonotypes (data not shown). These data suggested that infection-driven inflammation, not necessarily antigen re-encounter, is the driver of mTreg transition to xTregs. However, cognate antigen exposure may still be essential for the mTregs to mount an effective recall. At the numbers adoptively transferred, the PbT-II or OT-II mTregs remained undetected in unchallenged recipients (data not shown), and the transferred PbT-II mTregs undergoing homeostatic proliferation in RAG2KO mice maintained their Foxp3 expression (**Figure S9**). This suggests that infection-driven inflammation may be essential for the downregulation of Foxp3 in mTregs during recall responses.

Of note, the OT-II mTregs induced by *Lm-OVA*-infection also underwent robust recall expansion (indicated by Ki67 expression) and Foxp3 downregulation following re-exposure to *Lm-OVA* or *Py-OVA* infections (**Figure 4H-K**). This finding indicated that the transition of mTreg to xTregs during memory recall extends beyond malaria and is likely generally applicable to other infections and disease models.

### Transcriptional and epigenetic reprogramming underlies mTregs conversion to xTregs

To define the molecular and phenotypic changes driving mTreg-to-xTreg conversion, we performed single-cell transcriptomic and epigenetic analyses on mTregs and xTregs derived from *Pcc*-infected mice (**Figure 5A**). Unsupervised clustering revealed distinct but spatially proximate mTreg and xTreg populations, reflecting shared transcriptional features and likely developmental continuum between the cell subsets (**Figure 5B, Table S1**). Nevertheless, consistent with our previous findings presented in **Figure 4C**, the xTregs exhibited a noticeable downregulation of *Foxp3* transcripts, clearly demarcating them from mTregs (**Figure 5C**).

**Figure 5.**
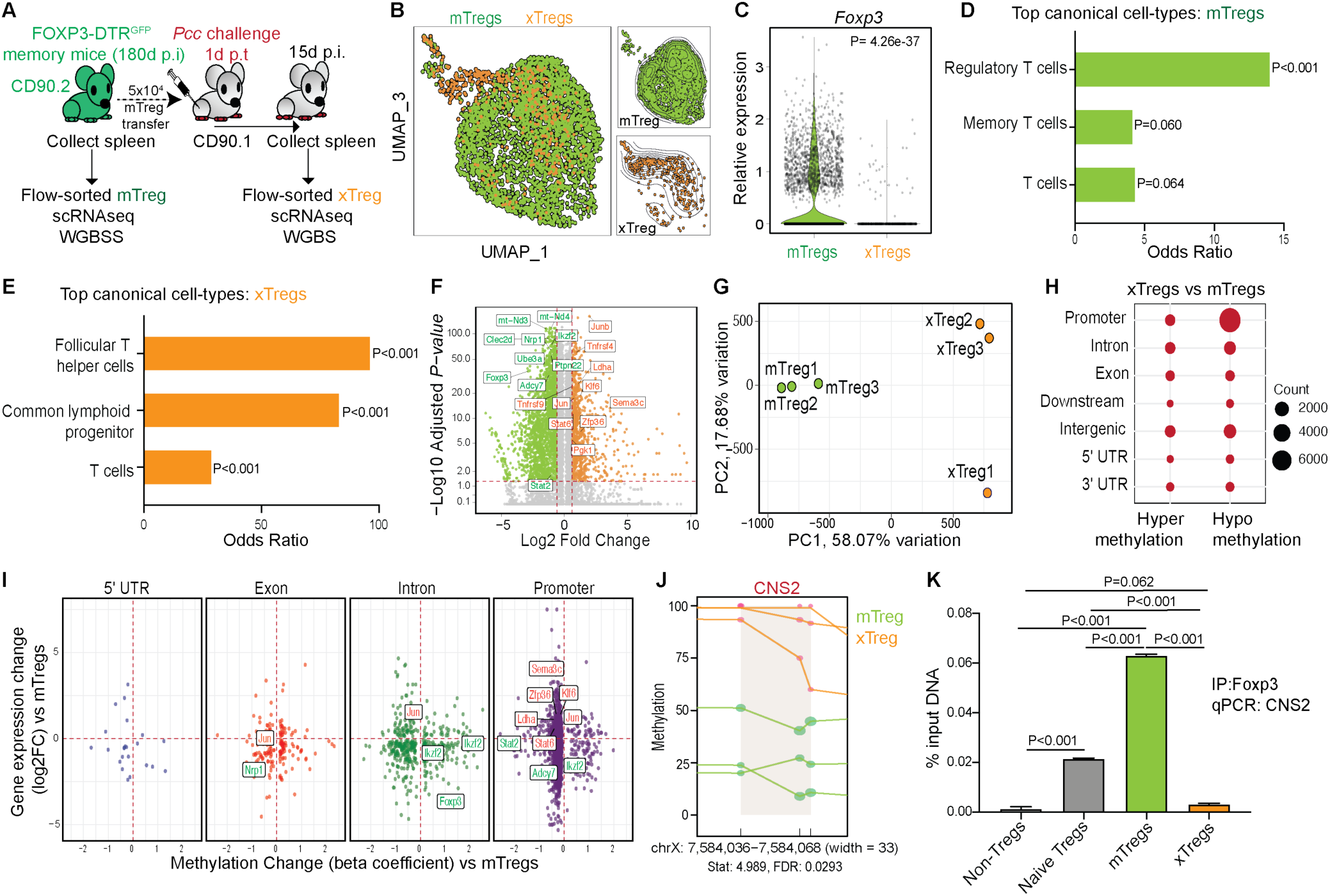
Transcriptomic and epigenetic differences between mTregs and xTregs. (A) Experimental scheme: 5x10^4^ flow-sorted splenic mTregs (CD90.2⁺) from *Pcc*-experienced FOXP3-DTR^GFP^ mice were transferred to congenically distinct B6 mice (CD90.1) followed by *Pcc* challenge (5x10^5^ iRBC/mouse). The molecular signatures of mTregs (from donors, 180d p.i.) and xTregs (recovered from recipients, 15d p.i.) were profiled using single-cell RNA sequencing (scRNAseq) and whole-genome bisulfite sequencing (WGBS) workflows; n=3. (B) Uniform Manifold Approximation and Projection (UMAP) of mTregs and xTregs based on their gene expression profiles (C) Violin plot depicting the expression of *Foxp3* in mTregs and xTregs. (D-E) Top cell lineage identities ascribed to mTregs (D) and xTregs (E) based on their transcriptomic signatures. Bar lengths represent the odds of transcriptional alignment with a canonical cell-type in mice; the numbers represent the adjusted *P*-value. (F) Volcano plot displaying the transcriptional differences between mTregs and xTregs, with top genes defining Treg and Tfh-like signatures marked. (G) Principal component analysis (PCA) plot illustrating the differences in genome-wide DNA methylation patterns in mTregs and xTregs. (H) Dot plots displaying the distribution of DNA methylation changes (units) in specific genomic regions of xTregs relative to mTregs. (I) Scatter plots of methylation versus gene expression changes in xTregs relative to mTregs, faceted by genomic regions. (J) DNA methylation profile of the CNS2 region in *Foxp3* in mTregs and xTregs. Dot size corresponds to relative read depth, each color-coded line connecting methylated loci represents an individual biological replicate (n =3 per cell type). (K) ChIP-qPCR analysis of Foxp3 association with *Foxp3* CNS2 in the indicated cell subsets. Data presented as mean ± SD normalized to input DNA from 1 of 2 separate experiments with 3 biological replicates, analyzed using one-way ANOVA with Tukey’s correction yielding the indicated *P*-values. Naïve Tregs (Foxp3^+^ CD4 T cells) or non-Tregs (Foxp3^-^ CD4 T cells) were derived, respectively, from naïve or *Pcc*-infected (180d p.i.) FOXP3-DTR^GFP^ mice.

To objectively define the cellular identity of xTregs, we performed unbiased computational alignment using Azimuth and Panglao databases for reference-based mapping and marker-based validation against the Tabula Muris reference atlas, identifying their closest canonical cell-type counterparts^47^. As expected, and serving as a validation of our approach, the mTregs obtained from *Pcc*-experienced mice aligned with canonical *Foxp3*^+^ Treg and memory T cell profiles (**Figure 5D**). In contrast, the xTregs exhibited a predominant Tfh-like signature, with a secondary common lymphoid progenitor (CLP)-like profile, suggestive of their transitional plasticity (**Figure 5E**). These assignments were supported by the upregulation of genes linked to immunoregulatory function (e.g., *Foxp3*, *Cd27*, *Ptpn22*), phenotype (e.g., *Foxp3*, *Ikzf2*, *Nrp1*), and persistence (e.g., *mt-Nd3*, *mt-Nd4*, *Stat2*) in mTregs, while the genes associated with Tfh differentiation (e.g., *Junb*, *Stat6*, *Tnfrsf4*, *Zfp36*) and effector functions (e.g., *Tnfrsf9*, *Ldha*, *Sema3c*, *Pgk1*) dominated in xTregs (**Figure 5F, Table S1**).

Epigenetic reprogramming is known to be the foundation of CD4 T cell plasticity^48, 49^. Therefore, we predicted that mTreg-to-xTreg conversion involved DNA methylation changes. Genome-wide methylation profiling of mTregs and xTregs revealed distinct clustering by principal component analysis (PCA) and identified 12,219 differentially methylated regions (adjusted *P* < 0.05) distributed across discrete genomic locations, underscoring their epigenetic divergence (**Figure 5G-H, Figure S10A, Table S2**). The exons, introns, and promoter regions of the various genes differentially expressed in memory, Treg, or Tfh cells showed distinct methylation patterns in the xTregs, suggesting that epigenetic alterations may be the basis of their transition from mTregs (**Figure 5I, Table S3**). Particularly significant among these changes was increased methylation of the Treg-specific enhancer, conserved non-coding sequence 2 (CNS2) within the *Foxp3* locus in xTregs (**Figure 5J**); hypermethylation of CNS2 is associated with inefficient Foxp3 binding, downregulation of *Foxp3* expression, and destabilization of the Treg transcriptional program^49, 50^. This observation was corroborated by chromatin immunoprecipitation-quantitative PCR (ChIP-qPCR) analysis showing that Foxp3 binding to CNS2 was significantly reduced in xTregs compared to mTregs, to the levels similar to that of in conventional (Foxp3^-^) non-Tregs (**Figure 5K**). Reciprocally, the Bcl6 promoter region exhibited reduced DNA methylation in xTregs (**Figure S10B**). Considering that Bcl6 is the master regulator of Tfh program in CD4 T cells, this suggested enhanced stability and function of the Tfh program in the xTregs^23, 51, 52, 53^. Together, the above data suggest that mTreg-to-xTreg conversion involves broad transcriptional changes and genome-wide epigenetic restructuring, extending beyond the isolated loci of their lineage-defining transcription factors.

### xTregs exhibit Tfh-like phenotypic and functional identity

Our transcriptional and epigenetic analyses predicted that xTregs would exhibit Tfh-like characteristics. Indeed, the polyclonal *Pcc*-specific xTregs showed near-complete Foxp3 downregulation while acquiring robust Bcl6 expression, providing experimental validation for this hypothesis (**Figure 6A-B**). This Tfh-lineage commitment was further evidenced by surface expression of CXCR5 and PD-1 (**Figure 6C**), which are signature markers of Tfh cells, and essential for germinal center homing and function^54^. Microscopy studies validated these findings, with the transferred mTregs localizing predominantly to the GCs in the spleen following *Pcc* infection in the recipient mice (**Figure 6D-E**). The xTregs formed clusters with B cells within the GCs, reminiscent of the characteristic Tfh-GC B cell interactions observed in the secondary lymphoid organs of *Plasmodium*-infected mice (**Figure 6F, Video S1**)^17^. The acquisition of Tfh-like phenotype also appeared consistent across different antigen specificities and infections: PbT-II or OT-II mTregs upregulated Bcl6 during recall induced by re-exposure to *Pcc* or *Lm-OVA* infections, respectively (**Figure S11**).

**Figure 6.**
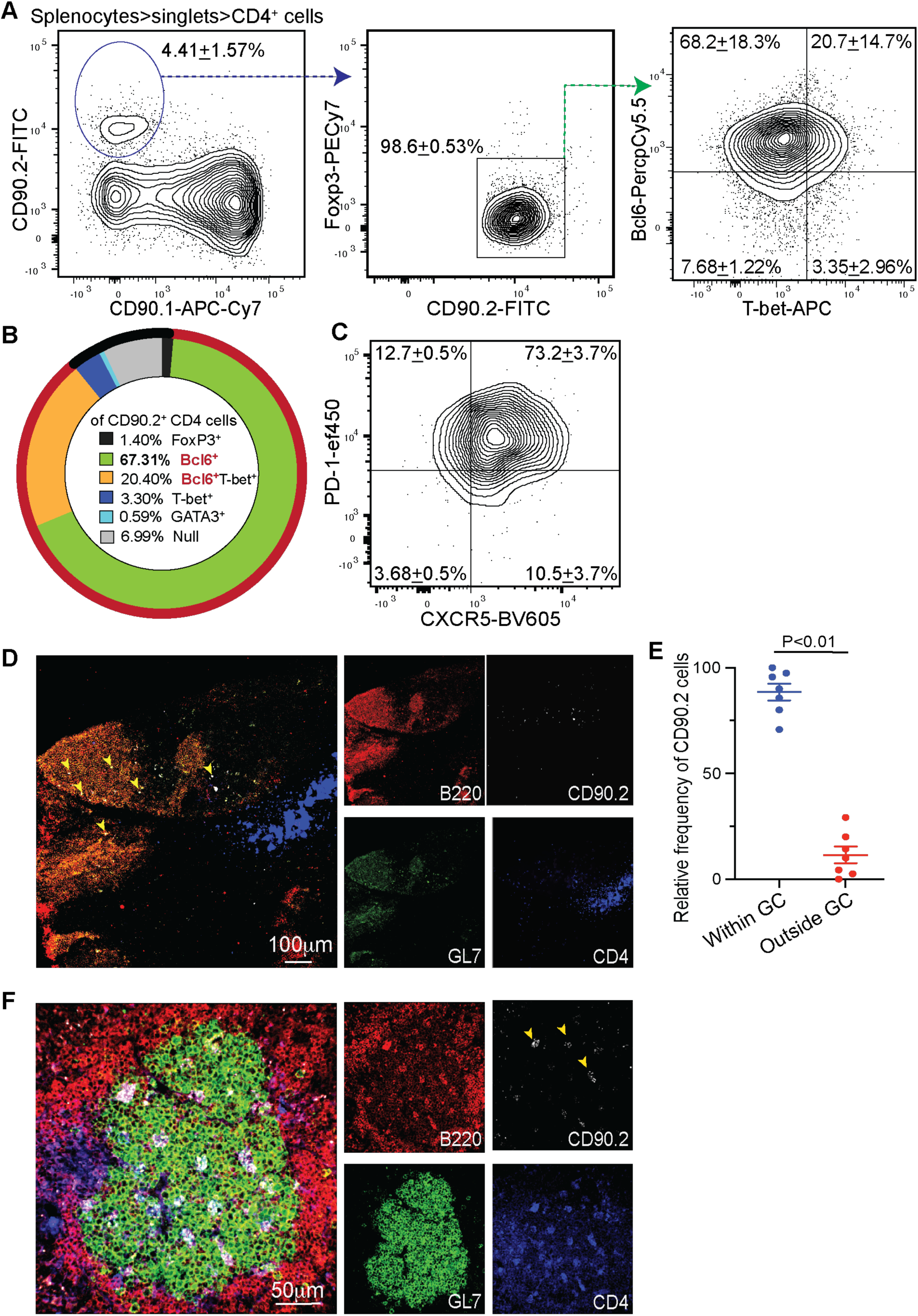
Memory Tregs transition to Tfh-like cells upon re-encountering *Plasmodium* infection. (A-C) As in Figure 5A, 5x10^4^ flow-sorted splenic mTregs (CD90.2⁺) from *Pcc*-experienced FOXP3-DTR^GFP^ mice were adoptively transferred to CD90.1⁺ B6 mice followed by *Pcc* challenge (5x10^5^ iRBC/mouse). Representative flow plots showing the frequency of the adoptively transferred cells expressing Foxp3, Bcl6, or T-bet in recipient spleens at 15d p.i. (B) Relative frequencies of the transferred cells expressing the indicated transcription factors. (C) Flow plot showing PD-1 and CXCR5 expression in the donor-derived cells obtained from recipient spleens. (A, C) Numbers inset indicate the gated cell frequencies, presented as mean ± SD from 1 of 3 identical experiments; n=3 recipients. (D) Representative pseudocolored confocal microscopy images showing localization of xTregs within the recipient spleen; yellow arrows indicate the transferred cells, 15d p.i. Data from 1 of 3 replicate separate experiments; n=3 recipients. (E) Scatter plot summarizing the relative proportions of xTregs located within the GC margins defined by GL7 expression. Data combined from 3 replicate experiments, presented as mean ± s.e.m., analyzed using t-test to yield the indicated *P*-value; each dot represents a microscopy field. (F) Representative pseudocolored confocal microscopy images showing xTreg clustering within GCs in recipient spleen at 15d p.i.; yellow arrows indicate xTreg clusters. Data from 1 of 3 replicate separate experiments; n=3 recipients.

Intriguingly, approximately 20% of the recalled xTregs co-expressed the Th1 associated transcription factor, T-bet, alongside Bcl6 (**Figure 6B**). A detailed analysis of T-bet and Bcl6 expression dynamics during mTreg-to-xTreg transition following *Pcc* infection indicated that the adoptively transferred T-bet^+^ Bcl6^-^ (Foxp3^+^) mTregs switched to the predominantly T-bet^-^ Bcl6^+^ Foxp3^-^ xTreg phenotype through an intermediary T-bet^int^ Bcl6^int^ stage (**Figure S12A-B**). The preservation of T-bet expression in xTregs suggested that they may retain some Th1-like properties. The xTregs also produced detectable IFN-γ following non-specific stimulation with calcium ionophores (**Figure S12C**). However, neutralizing IFN-γ in the TCRaKO recipients of mTregs did not significantly impair *Pcc* control (**Figure S12D-E**), suggesting that IFN-γ may be dispensable for mTreg-mediated protection from *Pcc* challenge. To test this more stringently, we generated FOXP3^βIFNγ^ (FOXP3-Cre^YFP^ crossed with IFNg^fl/fl^^55^) mice, in which the *Ifng* gene is ablated specifically in the Tregs. Transfer of FOXP3^βIFNγ^ mTregs protected TCRaKO recipients similar to wild-type mTreg transfers (**Figure S12F-G**). These data also align with the finding that IFN-γ produced by CD4 T cells have limited impact on the control of blood-stage malaria, particularly during recall^20, 55^. Together with our transcriptional and epigenetic analyses, these functional studies demonstrated that xTregs undergo comprehensive reprogramming to adopt a GC-homing Tfh phenotype, completing their transition from immunosuppressive cells to facilitators of protective immunity upon rechallenge.

### Bidirectional plasticity between Tregs and non-Tregs during memory formation

Recent evidence of Tfh cells acquiring a Treg phenotype during memory development^24^, complemented by our observations of mTregs converting to Tfh-like cells during recall, prompted us to investigate the origins of mTregs. Are mTregs derived solely from the expanded Treg population in a primary infection, as conventional memory development would dictate, or are they also derived from non-Tregs? To test this, we flow-sorted Foxp3^+^ or Foxp3^-^ CD4 T cells during the acute stage of *Pcc* infection (15d p.i.) and transferred them into congenically distinct, infection-matched hosts and mapped their fates during memory formation (**Figure S13A**). Strikingly, approximately 50% of the Foxp3^+^ cells downregulated their *Foxp3* expression, while ∼50% of the Foxp3^-^ cells upregulated *Foxp3* by 60d p.i. (**Figure S13B-C**). The interconversion between Treg and non-Treg states during memory development suggested that mTregs might arise from both conventional Tregs and reprogrammed non-Treg precursors, revealing an unappreciated fluidity in CD4 T cell memory populations. Only future lineage tracing studies of mTregs during its generation and recall through repeated antigen exposures will reveal the biological nature and significance of this plasticity.

### Tfh-like reprogramming essential is for mTreg-mediated protection against *Plasmodium*

To determine the functional significance of the acquisition of Tfh-like phenotype by mTregs, we generated mice that lack Bcl6 specifically in the *Foxp3*^+^ cells (FOXP3^ΔBcl6^), by cross-breeding FOXP3-Cre^YFP^ mice with Bcl6^fl/fl^ mice. A potential caveat of this mouse model, however, is the ablation of Bcl6^+^ Foxp3^+^ CD4^+^ follicular Tregs (Tfrs). Although a Tamoxifen-inducible Cre system would have overcome this limitation, Tamoxifen treatment has direct anti-malarial effects, precluding the application of such a model in this scenario^56, 57^. Considering that the proportion of Tfr cells is negligible within the mTregs compartment, we anticipated it to not be a confounding factor in the experiments with the FOXP3^ΔBcl6^ mice (**Figure S12A**).

We transferred congenically distinct mTregs from *Pcc*-experienced control FOXP3- Cre^YFP^ mice or FOXP3^βBcl6^ mice into naïve B6 recipients and challenged the latter with *Pcc* (**Figure 7A**). While the FOXP3-Cre^YFP^ mTregs boosted splenic activated CD4 T cell, Tfh cell, B cell, GC B cell, and GC plasmablast numbers, the responses in FOXP3^βBcl6^ mTreg recipients resembled the no-transfer controls (**Figure 7B-F**). Corroborating these findings, FOXP3^βBcl6^ mTregs induced significantly diminished GC reactions in the spleen (**Figure 7G-J**). Microscopy analyses also revealed that while control xTregs localized within the GCs and established noticeable B cell clusters there, the absence of Bcl6 prevented the recalled mTregs from establishing such clusters in the recipient GCs or display any detectable engagement with the GC B cells (**Figure 7K-L, Videos S2, S3**). The functional consequences of Bcl6 ablation in mTregs were equally striking: although both FOXP3-Cre^YFP^ and FOXP3^ΔBcl6^ mTreg transfers helped mount anti-malarial IgM responses, only FOXP3-Cre^YFP^ mTregs recipients developed robust IgG titers (**Figure 7M-N**). This is consistent with the role of Bcl6 and GC Tfh cells in facilitating class-switch recombination during GC reaction^58, 59^. Corroborating these findings, unlike the FOXP3-Cre^YFP^ mTregs, FOXP3^βBcl6^ mTregs were unable to transfer protection against *Pcc* challenge in TCRaKO mice (**Figure 7O**).

**Figure 7.**
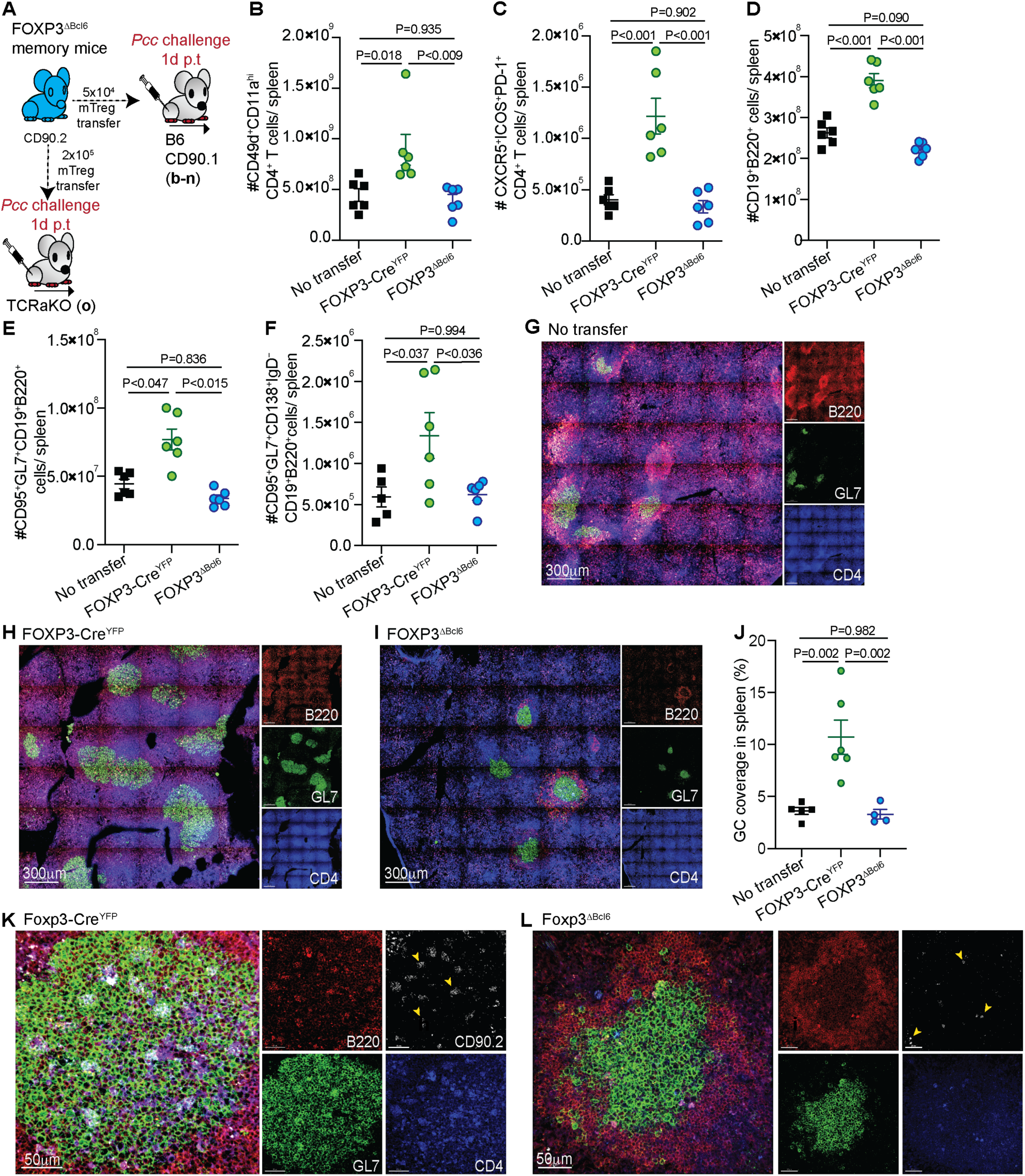
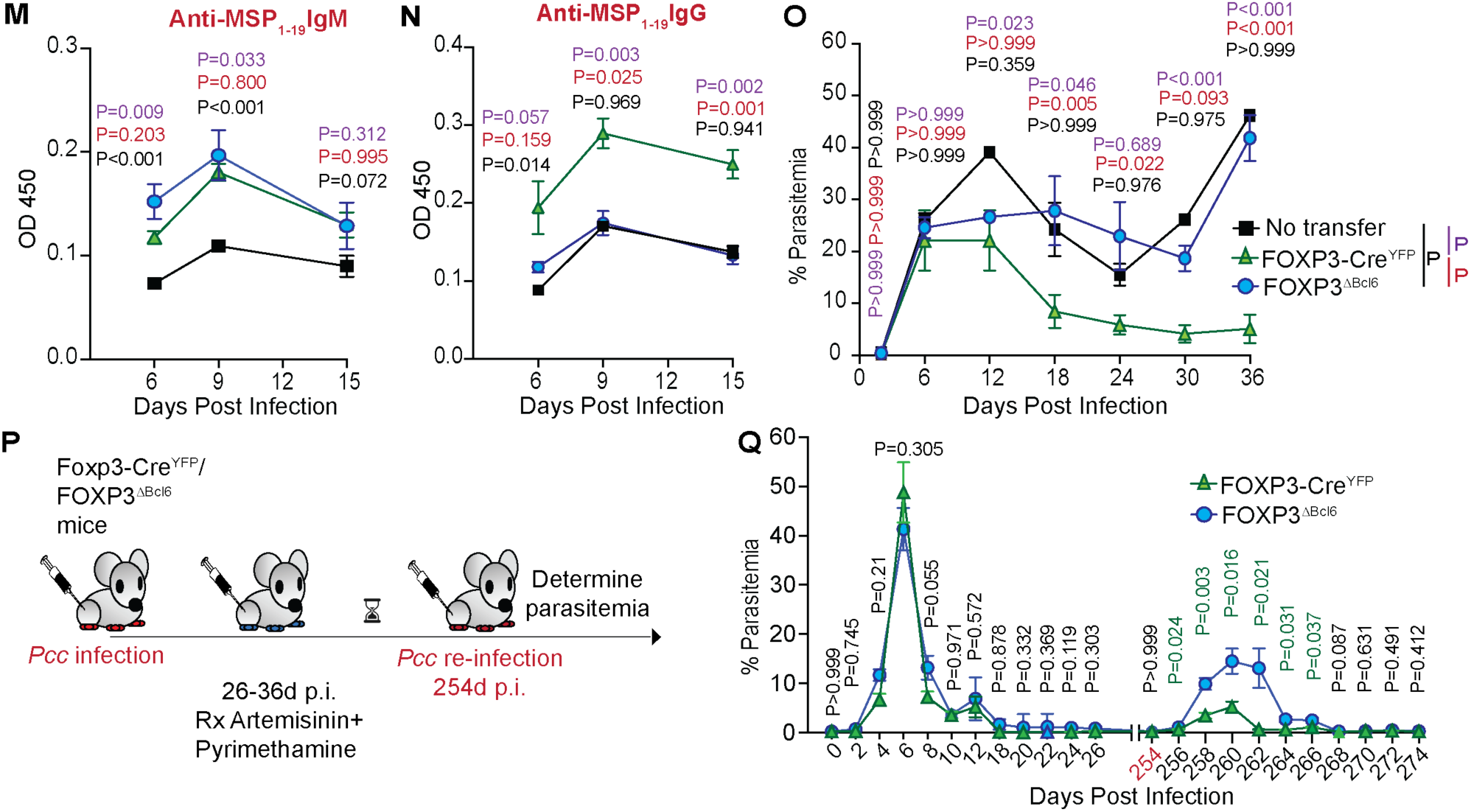
Bcl6 essential for mTreg-mediated protection from *Plasmodium* challenge. (A) mTregs derived from *Pcc*-experienced FOXP3^βBcl6^ or control FOXP3-Cre^YFP^ mice were adoptively transferred to TCRaKO or congenically distinct B6 recipients, which were then challenged with *Pcc* (5x10^5^ iRBC/mouse). B6 mice receiving no transfer served as additional controls. (B-F) Scatter plots indicating the total numbers of *Plasmodium*-specific CD4 T cells (B), Tfh cells (C), B cells (D), GC B cells (E), or GC plasmablasts (F) in recipient spleens at 15d p.i.; combined data from 2 experiments presented as mean ± s.e.m, each dot represents a recipient. (G-J) Representative pseudocolored confocal images showing splenic GC reaction in the recipients of no transfer (G), FOXP3-Cre^YFP^ (H) or FOXP3^βBcl6^ (I) mTreg transfers, 15d p.i. Representative data from 1 of 2 experiments with 3 mice/group. Scatter plots summarize splenic GC coverage in these recipients; representative data presented from 1 of 2 experiments as mean ± SD; each dot represents a recipient (J). (K and L) Representative pseudocolored confocal microscopy images showing FOXP3-Cre^YFP^ (K) or FOXP3^βBcl6^ (L) xTreg localization relative to the GCs in the recipient spleen; arrow heads indicate xTreg clusters, represents ≥15 fields observed in a total of three separate mice from two separate experiments. (M and N) Anti-MSP_1-19_ IgM (M) or IgG (N) serum antibody titers in the no transfer, FOXP3^βBcl6^, and FOXP3-Cre^YFP^ recipient mice following *Pcc* infection, at the indicated time points p.i. Representative data from 1 of 2 experiments, presented as mean ± SD, analyzed using 2-way ANOVA with Tukey’s correction, yielding the indicated *P*-values. (O) Temporal kinetics of parasitemia in TCRaKO, with no, or mTreg transfers from FOXP3-Cre^YFP^ or FOXP3^βBcl6^ donors, challenged with *Pcc* iRBCs. Representative data from 1 of 3 replicate experiments, presented as mean ± SD, analyzed using two-way ANOVA with Sidak correction, yielding the indicated *P*-values. (P) Experimental scheme: FOXP3-Cre^YFP^ or FOXP3^βBcl6^ mice infected with *Pcc* (5x10^5^ iRBC/mouse) were drug-cured and re-challenged with *Pcc* (2x10^6^ iRBC/mouse) at 254d p.i. (Q) Temporal kinetics of parasitemia; data from 2 replicate experiments, presented as mean ± s.e.m, analyzed using 2-way ANOVA with Sidak correction, yielding the indicated *P*-values.

To model the malaria reinfection cycle observed in human patients residing in malaria-endemic areas, we next examined the ability of *Pcc*-experienced FOXP3-Cre^YFP^ or FOXP3^βBcl6^ mice to control *Pcc* reinfection (**Figure 7P**). While both FOXP3-Cre^YFP^ and FOXP3^βBcl6^ mice controlled the primary *Pcc* infection to a similar extent, FOXP3^βBcl6^ mice exhibited a significant defect in limiting the secondary challenge with *Pcc* (**Figure 7Q**). A 2-to-3-fold increase in parasitemia, as in the case above, can elevate an asymptomatic and sub-clinical case of malaria to severe clinical disease accompanied by multi-organ failure and death in humans^60^.

Together, these findings supported the notion that mTreg transition to Tfh-like cells may be essential to offer protection from repeat infections with *Plasmodium*. Our work establishes a direct mechanistic link between the transcriptional reprogramming and the functional transition of the immunosuppressive mTreg population to protective xTregs, offering new insights into how memory Tregs adapt to recurrent infections.

## Discussion

Our study reveals that memory Tregs generated during *Plasmodium* infection undergo functional reprogramming upon rechallenge, transitioning from immunosuppressive cells to Tfh-like effectors that promote germinal center responses and protective immunity. This discovery fundamentally challenges three immunological dogmas: that Treg memory exists to reinforce suppression, memory lineages remain strictly segregated, and that recurrent infections merely reactivate static cell populations. The implications of these findings extend beyond malaria, suggesting a radical reinterpretation of how immune memory balances regulation and protection.

We consider the existing paradigm of mTregs becoming increasingly suppressive with each antigen encounter to be evolutionarily paradoxical. Why preserve a system that recalls suppression when rapid effector responses are most needed? Although clonal expansion of small populations of antigen-specific Tregs may be necessary in certain contexts such as in maintaining tolerance to repeated exposures to pregnancy-associated alloantigens, food antigens, or commensal microbiota-derived antigens, is this applicable to recurrent infections? If classical memory responses evolved to avoid the inflammatory costs of primary infections, concurrent reactivation of suppression would seem counterproductive to that cause. Our data, at least partly resolve this paradox by suggesting that mTregs may not be static enforcers of suppression, but rather dynamic sensors of the inflammatory context. We propose that, in the context of infections, during primary exposure, the suppressive functions of Tregs would prevent immunopathology. During recurrence of such infections, epigenetic reprogramming would convert mTregs into Tfh-like cells (xTregs) that accelerate humoral memory and pathogen clearance in an antigen-specific manner^61^.

However, this perception is complicated by our finding that not all mTregs arise from *bona fide* Tregs. This also aligns with the recent report of Tfh cells acquiring a Treg phenotype during the dissolution of the GCs in the lymphoid organs during the memory phase, after immunization^24^. While the timing of assessment during the progression to immunological memory may dictate the extent to which Tregs and non-Tregs contribute to the existing population of mTregs, it raises an important question: if antigen-specific recall is the intent of immunological memory, why would non-Tregs take on the phenotype of Tregs during the memory stage? One consideration is that Foxp3 expression is associated with increased IL-2Rα expression in T cells^62^. Enhanced IL-2Rα expression would allow Th cells to be maintained in the diminishing IL-2 environments associated with the post-infection and immunization period. Of note, the majority of the Th cell subsets outside of Tregs are notoriously short-lived and acquisition of Foxp3 expression may serve as a solution for that insufficiency^63^.

A related question is whether the recalled xTreg population would undergo attrition after serving their effector purpose, or might they revert to Foxp3^+^ states to rejoin the (presumably) longer-lasting memory pool? Our snapshot likely captures just one phase of the dynamic process of memory development and recall. The fact that Tfh cells can re-acquire Foxp3 expression post-GC resolution implies that the cell subset we consider mTregs may include various transitional states along an effector-regulatory continuum. Our single cell analysis displaying transcriptional heterogeneity within the mTregs and xTregs supports this premise. While only future lineage tracing studies with single-cell resolution across multiple infection cycles would provide a definitive answer to this question, we favor the prediction that xTregs re-express Foxp3, allowing them to “reset” for future recall. This cyclical adaptation model finds support in observations that conventional effector T cells can transiently upregulate Foxp3 for metabolic benefits^64^. This fluidity also aligns with growing recognition of CD4 T cell plasticity across discrete immunological contexts^22^. Our own discovery of the molecular basis of the conversion of mTregs to Tfh-like xTreg via the epigenetic switch from Foxp3 to Bcl6 expression provides mechanistic credibility to this paradigm.

Our findings also demand a fundamental reconsideration of Treg memory biology. Rather than serving as dedicated suppressors, mTregs emerge as multipotent adaptors that toggle between regulatory and effector states according to contextual demands- a plasticity we propose enables the immune system to retain antigen-experienced cells in cytokine-limited environments without the burden of constitutive suppression. This represents an optimal strategy for recurrent infections, where rigid lineage commitment, especially with Tregs, may compromise protective immunity. The clinical and therapeutic implications of such findings may be profound. While harnessing this adaptability could revolutionize interventions for recurrent infections, autoimmunity, and cancer immunotherapy, it will require careful navigation of the context and time-dependent functional dichotomy of mTregs during antigen persistence or re-exposure. The frequency of pre-existing mTregs could also serve as predictive biomarkers for a better prognosis, as in the case of human malaria. Ultimately, our malaria studies challenge the principle that mTregs are stable enforcers of tolerance, revealing them instead as possible ‘immunological shapeshifters’, which are equipped to persist through Foxp3-mediated survival mechanisms, yet poised to provide help when recurrent challenges demand it.

## Methods

### Malian and U.S. blood donors

The Ethics Committee of the Faculty of Medicine, Pharmacy and Dentistry at the University of Sciences, Technique, and Technology of Bamako and the National Institute of Allergy and Infectious Disease of the National Institutes of Health (NIAID/NIH) Institutional Review Board approved the study in Mali (protocol no. 11-I-N126). The field study, described in detail elsewhere, was conducted in Kalifabougou, Mali, where malaria transmission occurs during the 6-month rainy season^11^. Blood samples were obtained from study participants at two time points: 1) at their healthy uninfected baseline (before the malaria season) at least 180 days after being treated for acute febrile malaria, and 2) during their first febrile malaria episode of the ensuing malaria transmission season. Malaria cases were defined as an axillary temperature >37.5°C and >2500 asexual *P. falciparum* parasites/µl of blood, with no other cause of fever identified by medical history and physical examination. Blood samples of healthy U.S. study participants were obtained from the NIH blood bank for research use acquired via the protocol approved by the NIH Institutional Review Board (protocol no. 99-CC-0168). Written informed consent was obtained from all study participants or from the parents or guardians of participating children. All PBMC samples were analyzed at the NIAID/NIH.

### Mice and pathogens

C57BL/6, FOXP3-DTR^GFP^ (B6.129(Cg)-*Foxp3^tm3^*^(*DTR*/*GFP*)*Ayr*^/J)^42^, FOXP3-Cre^YFP^ (B6.129(Cg)-*Foxp3^tm4(YFP/icre)Ayr^*/J)^65^, Bcl6^fl/fl^ (B6.129S(FVB)-*Bcl6^tm1.1Dent^*/J)^66^ and TCRaKO mice were purchased from the Jackson Laboratories. IFNγ^fl/fl^, PbT-II and OT-II mice were gifts from Drs. John Harty (University of Iowa, USA), William Heath (The Peter Doherty Institute for Infection and Immunity, Australia), and Wendy Watford (University of Georgia, USA), respectively. The animals were treated and handled in accordance with guidelines established by UGA Institutional Animal Care and Use Committees and housed with appropriate biosafety containment at the University of Georgia (UGA) animal care units. All mice were infected at 6-12 weeks of age.

*P. chabaudi chabaudi* AS (*Pcc*) parasite was gifted by Dr. Julie Moore (University of Florida), *P. yoelii* 17XNL (*Py*) and *P. berghei* ANKA (*Pb*) were obtained from BEI resources, *Py-OVA* and *Pb-OVA* were generated in our lab as described before^67, 68^. *L. monocytogenes* (*Lm* ActA^-^) and *L. monocytogenes-OVA* (*Lm-OVA*) were gifted by Dr. John Harty. All infections in mice were initiated with 5-20 x 10^5^ *Pcc*, *Py*, *Pb*, *Py-OVA* or *Pb-OVA* infected erythrocytes (iRBCs) inoculated intravenously (i.v), 1 x 10^7^ colony-forming units (CFU) of *Lm* or *Lm-OVA,* i.v., or 1×10^3^ or 5×10^2^ plaque-forming units (PFU) of influenza A virus (X31 or PR8 strains) intranasally (i.n.).

### Flow cytometry

For human studies, PBMCs were isolated and stained, and FACS analysis was performed as described previously^69^. For mouse studies, single-cell suspensions of cells obtained from spleen or blood were stained mouse antibodies as described before^13^. All reagents were purchased from BioLegend, BD Biosciences or eBiosciences. For human studies, human CD3 (clone UCHT1), CD4 (clone SK3), CD45RO (clone UCHL1), CD45RA (clone HI100), CXCR5 (clone MU5UBEE), Foxp3 (clone 259D), Helios (clone 22F6), and CTLA-4 (clone 14D3) were employed. In the case of mice, antibodies against CD4 (clone GK1.5), CD8 (clone 53-6.7), CD49d (clone R1-2), CD11a (clone M17/4), CD27 (clone LG.3A10), CD127 (clone A7R34), ICOS (clone C398.4A), CD62L (clone MEL-14), CD69 (clone H1.2F3), KLRG1 (clone 2F1/KLRG1), CD25 (clone PC61), Nrp1 (clone V46-1954), PD-1 (clone RMP1-30 or J43), CXCR5 (clone L138D7), CD44 (clone IM7), CD90.1 (clone OX-7), CD90.2 (30-H 12), CD19 (clone 1D3), GL-7 (clone GL7), CD95 (clone SA367H8), CD138 (clone 281-2), IgD (clone 11-26c.2a), B220 (clone RA3-6B2), TCRvα2 (clone B20.1), TCRvβ12 (clone MR11-1), TCRvβ5.1 (clone MR9-4) or Ki67 (clone 16A8) were used for surface staining. For intracellular stains, antibodies against CTLA-4 (clone U10-4B9), T-bet (clone AB10), GATA3 (clone TWAJ), Bcl6 (clone 7D1), Foxp3 (clone 3G3 or MF23) or RORγt (clone B2D), BrdU (clone BU20A), and IFN-γ (clone XMG1.2) were employed. Foxp3 staining kit (eBiosciences) was used for intracellular staining according to the manufacturer’s instructions. Multicolor flow cytometry was performed on Agilent Novocyte Quanteon or CytoFLEX analyzers and results were analyzed with FlowJo software (Tree Star).

### Cell Sorting

Sorting of murine splenic lymphocytes from GFP or YFP reporter mice was performed as previously reported^70^. Briefly, spleens were homogenized in Ammonium-Chloride-Potassium (ACK) lysis buffer, filtered through 70 µm strainer and washed in Magnetic Activated Cell Sorting (MACS, Miltenyi) buffer. Single-cell suspensions of the splenocytes were then enriched for CD4 T cells (Miltenyi MACS or STEMCELL EasyEight^TM^ EasySep^TM^) before incubating with one or more of anti-mouse antibodies to the cell surface markers CD4, CD44, CD90.2, TCRvα2, TCRvβ12, or TCRvβ5.1 as applicable for 10-20 min at 4°C. Cells were then washed and resuspended in MACS buffer and flow-sorted on Modular Flow Cytometer (MoFlo, Beckman Coulter), Aurora CS (Cytek) or FACSAria II cell sorter (BD Biosciences) under sterile conditions. Naive Tregs (polyclonal) were defined as GFP^+^ CD4 T cells or YFP^+^ CD4 T cells from FOXP3-DTR^GFP^ or FOXP3-Cre^YFP^ mice respectively; mTregs were defined as GFP^+^ CD44^+^ CD4 T cells from FOXP3-DTR^GFP^ mice. Non-Tregs were defined as GFP^-^CD4^+^ T cells. PbT-II naïve or memory Tregs were identified as GFP^+^ TCRvα2^+^ TCRvβ12^+^ CD4^+^ T cells or GFP^+^ TCRvα2^+^ TCRvβ12^+^ CD44^+^ CD4^+^ T cells from PbT-II-FOXP3-DTR^GFP^ mice, respectively; OT-II naïve or memory Tregs were defined as GFP^+^ TCRvβ5.1^+^ CD4^+^ T cells or GFP^+^ TCRvβ5.1^+^ CD44^+^ CD4^+^ T cells from OT-II-FOXP3-DTR^GFP^ mice.

### *In vitro* Suppression Assay

Standard *in vitro* Treg suppression assays used for assessing Treg function were modified from established protocols described in detail before^71^. In short, 5x10^4^ CellTrace Violet (CTV)-labeled OT-I CD8 T cells were co-cultured with 5x10^4^ LPS-activated dendritic cells isolated as described before^72^ and pulsed with chicken OVA_257-264_ (SIINFEKL peptide) antigens, either in the absence or presence of serially diluted mTregs in RPMI media (Sigma-Aldrich) reconstituted with 10% FBS (Sigma-Aldrich) in a 96-well plate, at 37°C, for 72 hours. Naive Tregs served as controls. Frequency of OT-I CD8 T cell proliferation was assessed for CTV dilution by flow cytometry at the end of incubation.

### *In vitro* Germinal Center Reaction

The *in vitro* germinal center (GC) reaction was established as described in detail elsewhere^73^. In short, 5x10^4^ flow-sorted splenic naïve GC B cells (CD19^+^ B220^+^) and Tfh cells (CXCR5^+^ PD-1^+^ CD4^+^) were co-cultured with 10 µg/ml each of anti-CD3 and anti-IgM antibodies in IMDM media (Gibco) supplemented with 10%FBS (Sigma-Aldrich) in a 96-well plate, at 37°C, for 72-96 hours. Serial dilutions of naïve Tregs or mTregs were added to these cultures. The frequencies of CD95^+^ GL7^+^ B220^+^ CD19^+^ GC B cells were examined by flow cytometry at the end of incubation.

### Adoptive transfer

Adoptive transfers of murine lymphocytes were performed as we have previously described^74^. Briefly, FACS-purified lymphocytes were washed and resuspended in incomplete RPMI media (Sigma-Aldrich). Appropriate number of cells were then adoptively transferred to naïve or *Pcc-* experienced recipient mice by retro-orbital injections using hypodermic needles.

### Parasite kinetics

Parasitemia was assessed by flow cytometry as previously described in detail^13^. Briefly, whole blood samples were collected from the tail-end of infected mice, stained with Hoechst 33342 (Sigma), anti-Ter119-PE, and anti-CD45-APC antibodies constituted in phosphate buffer saline (1xPBS, Gibco). Data were acquired on Agilent NovoCyte Quanteon4025 and analyzed using FlowJo V10 (BD Life Sciences).

### Plaque assay

Plaque assays were performed as previously described in detail^75^. Briefly, lungs from IAV-exposed mice were lysed at the experimental endpoint in 1 ml MEM (Gibco) with 1 μg/ml TPCK-treated trypsin (Thermo Fisher). Confluent monolayers of Madin-Darby canine kidney (MDCK) cells were incubated with 10-fold serially diluted 10% lung tissue homogenate for 1h at 37°C. The inoculates were removed and the MDCK cells were washed with 1xPBS. Monolayers were then overlaid with MEM containing 1.2% Avicel microcrystalline cellulose (FMC BioPolymer), 20 mM HEPES, 2 mM L-glutamine, 1.5g/L NaHCO_3_, and 1 μg/ml TPCK-trypsin. At 72h of incubation, the MDCK monolayers were fixed with cold methanol and acetone (60:40), stained with crystal violet, visualized, and the plaques enumerated to extrapolate the plaque forming units per lung.

### Intracellular Cytokine Staining of IFN−γ

Assessment of intracellular IFN-γ production was performed as previously described^74^. Briefly, cells were cultured in RPMI media (Sigma-Aldrich) with 10% FCS, penicillin, streptomycin, L-glutamate, HEPES, B-mercaptoethanol, and gentamycin. After restimulation with phorbol myristate acetate (PMA) and Ionomycin (BioLegend) for 4h at 37°C, in the presence of Brefeldin A (BioLegend), the cells were stained for surface markers and fixed overnight at 4°C. Cells were then permeabilized using eBioscience PermWash buffer and stained with anti-mouse IFN-γ antibodies. Stained cells were then washed and resuspended in FACS buffer for flow cytometry.

### Therapeutic regimen

The following reagents along with the appropriate IgG controls or diluent were used: anti-IFN-γ monoclonal antibody (mAb) (catalog no. BE0055, BioXcell): 500 μg per mouse, i.p.; DT (Sigma-Aldrich): 1 μg per mouse, i.p.; Artemisinin (AT): 10 mg/kg bodyweight, i.p.; Pyrimethamine: 1.5 mg/kg i.p.; BrdU: 2 mg/kg i.p. followed by 0.8 mg/ml in drinking water.

### Microscopy

Mice spleen sections mice were fixed, stained, and imaged using the Zeiss LSM 880 laser scanning confocal microscope as described in detail previously^13^. In short, whole spleen was excised and fixed in 4% paraformaldehyde overnight. Spleen was then soaked in 15% sucrose solution for 2-4 h before it was transferred into 30% sucrose solution for an overnight incubation at 4°C. Spleen was then embedded in an O.C.T. compound (Scigen Scientific, Inc.) for a flash freezing in liquid nitrogen before storing at -80°C freezer. For staining, microscope slides of cryo-sectioned spleen were fixed in 100% acetone at -20°C for 30 min before rehydration at RT for 10 min. The cryo-sections were then blocked with OneBlock EIA (Prometheus) for 30 min at RT and stained for overnight at 4°C with direct fluorochrome-conjugated antibodies to CD4 (clone GK1.5), B220 (clone RA3-6B2), GL-7 (clone GL7) and/or CD90.2 (clone 30-H12) (BioLegend) constituted in 1% BSA/PBS using 1:50 or 100 dilutions. Cryo-sections were washed thrice and mounted in VectaShield antifade mounting medium (Vecta laboratories, Inc) for visualization using Zeiss LSM 880 confocal microscopy. Microscope images were analyzed and GC volumes determined using the ZEN (v2.3) and Imaris (V10.2) imaging software. GC volumes were determined using the ZEN 2.3 analysis software and the percentage GC coverage calculated by dividing the GC volume by splenic section volume.

### ELISA

MSP_1-19_-specific antibodies in sera were detected by ELISA as we have previously described in detail^13^. Briefly, serial dilutions of MSP_1-19_ protein were coated in triplicate in 96-well format in coating buffer (pH 9.0), overnight at 4°C, washed thrice with 0.05% Tween 20 in PBS (PBS-T), blocked for 120 min with 1% BSA/PBS, and incubated with serially diluted murine serum for 60 min at 37°C, washed five times with PBS-T, and then probed with peroxidase conjugated goat anti-mouse IgG HRP (1.0 mg/ml) or IgM HRP (0.8 mg/ml) as applicable, diluted in 1% BSA/PBS buffer, and incubated for 60 min at RT. Plates were then washed three times with PBS-T. All ELISAs were developed using a tetramethylbenzidine liquid substrate system (Sigma-Aldrich), stopped with 2 N sulfuric acid, and then read at 450 nm using an ELISA microplate reader (Molecular Devices SpectraMax iD3).

### Transcriptomic analysis

Generation and sequencing of single-cell RNA libraries: We prepared single-cell RNA-seq libraries in a similar manner as previously reported^76^. Briefly, mTregs and xTregs were enriched by flow-sorting. 10x Genomics’ Chromium Single Cell 3’ Gene Expression Technology (version 3) was used to generate single cell RNA-sequencing libraries of flow-sorted mTreg and xTregs. To this end, we added the sorted cells into independent wells of a Chromium Chip B containing the reverse transcription (RT) master mix. Next, we loaded the Chromium Chip B containing the cells, partitioning oil, and the Single Cell 3ʹ v3 Gel Beads into the 10x Genomics’ Chromium Controller for Gel Bead-in Emulsions (GEM) generation and cDNA barcoding. We then performed post-GEM-RT cleanup, cDNA amplification, and 3’ gene expression library construction as per the Chromium Single Cell 3ʹ Reagent Kits v3 User Guide (CG000183_ChromiumSingleCell3 v3_UG_Rev-A). We sequenced the cDNA libraries at the Georgia Genomics and Bioinformatics Center on an Illumina NextSeq 2000 platform using the run parameters specified in the Chromium Single Cell 3ʹ Reagent Kits v3 User Guide. Alignment of the single-cell RNA sequencing data to the *Mus musculus* reference genome: We performed quality control checks on the raw sequencing data using FASTQC and trimmed the reads using FASTX-Toolkit (version 0.0.14) to correspond to the appropriate R1 and R2 read lengths for 10x Genomics’ 3’ gene expression technology. Next, we aligned the sequencing data to the *Mus musculus* reference genome. To this end, we first generated a genome index using the *genomeGenerate* function in STAR (version 2.7.10a)^77^. We used the *M. musculus* genomic DNA assembly (GRCm39, Ensembl release 108), its corresponding gene annotation (gtf) file and the --sjdbOverhang parameter set to 90 for this task. We then used the single cell RNA-sequencing workflow (STARsolo) within STAR to map, demultiplex, and quantify gene expression^77^. We specified the following parameters to complete this task: --soloType CB_UMI_Simple --soloCBstart 1 --soloCBlen 16 --soloUMIstart 17 --soloUMIlen 12 -- soloBarcodeReadLength 1 --clipAdapterType CellRanger4 --soloCBmatchWLtype 1MM_multi_Nbase_pseudocounts --soloUMIfiltering MultiGeneUMI_CR --soloUMIdedup 1MM_CR --soloMultiMappers Unique Uniform Rescue PropUnique EM --soloStrand Forward -- soloFeatures Gene GeneFull SJ GeneFull_ExonOverIntron GeneFull_Ex50pAS --soloCellFilter EmptyDrops_CR --soloCellReadStats Standard.

Processing of single-cell RNA sequencing cell-gene count matrices: We imported the ‘GeneFull’ unfiltered (raw) matrix, features, and barcode files generated from STAR into R (version 4.3.2) using DropletUtils’ read10xCounts function^78^. We then used DropletUtils’ emptyDrops function with the lower argument set to 100 to distinguish GEMs containing cells from GEMs containing ambient RNA. We next performed quality control on the mTreg and xTreg datasets. We removed cells with fewer than 100 detected genes and excluded cells with mitochondrial gene content exceeding 35%. We also filtered out genes with total expression counts less than or equal to 2 across all cells. We next processed the datasets to identify and remove multiplets. To do this, we converted the data to Seurat objects and generated log-normalized gene expression values using the NormalizeData function in the Seurat R package^79^. We then performed multiplet detection using scDblFinder (version 1.20.2), retaining only cells classified as singlets and removing those identified as doublets. We annotated the data using SingleR, with bulk mouse RNA-seq data from sorted cell populations as the reference^80, 81^. We also performed unsupervised clustering of the data using a graph-based clustering approach as part of the Seurat R package^79^. We used the reference-based cell identification output to manually curate the unsupervised clustering outputs. Differential gene expression analysis and gene set enrichment analysis: We identified genes displaying differential expressions in mTreg and xTreg subpopulations (i.e., clusters) using Seurat’s FindAllMarkers function with the MAST parameter selected^82^.We considered genes as markers of a subpopulation if they were differentially expressed versus the other subpopulations and had a Bonferroni adjusted *P*-value <0.05. To identify genes differentially expressed between mTregs and xTregs, we first integrated the data using Seurat’s IntegrateData function. Next, we used Seurat’s FindMarkers function with the MAST parameter selected to perform differential expression testing^82^. We considered genes as differentially expressed if they had a Bonferroni-adjusted *P*-value <0.05 and a log₂ fold change greater than 0.585 or less than –0.585. We used the genes identified as differentially expressed (Avg. FC >1.5) as input for gene set enrichment analyses^83, 84^.

### Assessment of *FOXP3* expression in Tregs derived from human study participants

We downloaded the processed Seurat object (rna_tcr_malaria_specific.rds) from Nideffer *et al.* containing whole blood single-cell RNA-seq data from patients at various time points, including before and after repeated malaria^21^. We subsetted the Seurat object to assess *FOXP3* expression in resting CD4 T cells (stim = “unstimulated) identified as Tregs (cell_type = “Treg”, clone_family = “6”) in samples derived from patients before and after a confirmed malaria episode (sample = “3410_T3”, “3410_T4”, “3178_S7”, “3178_S8”, “3149_S3”, “3149_S4”, “3394_T1”, “3394_T2”, “3481_T1”, “3481_T2”, “3410_T1”, “3410_T2”, “3178_S5”, “3178_S6”, “3149_S1”, “3149_S2”). From these cells, we assessed Treg clones that displayed *FOXP3* expression (mean expression > 0) pre- and post-clinical episodes of malaria, and of which expanded after the infection. We used paired Wilcoxon signed-rank tests to compare mean *FOXP3* expression pre- and post-malaria within the TCR-restricted clones.

### DNA methylation analysis

Generation and sequencing for DNA methylation profiling at single-nucleotide resolution: Reduced-representation bisulfite sequencing (RRBS) libraries from mTregs (n = 3 biological replicates) were prepared using the Zymo-Seq RRBS Library Kit workflow (Cat. D5460, Zymo Research) according to the manufacturer’s protocol. Briefly, 500 ng of genomic DNA from each sample was used for MsP digestion, adapter ligation and gap filling, bisulfite conversion, and index primer (Zymo-Seq UDI Primer Set., Cat. D3008) amplification. Library quality control was performed on an Agilent 2100 TapeStation. Libraries were sequenced on an Illumina NextSeq2000 instrument (50 bp PE reads) with a PhiX control at 20% concentration of the total libraries.

Whole-genome bisulfite sequencing (WGBS) libraries from xTregs (n = 3 biological replicates) were prepared using a protocol adapted from Raine *et al.* 2022^85^. Briefly, 1.3 ng of genomic DNA from each sample was bisulfite converted using EZ DNA Methylation-Lightning Kit workflow (Cat. D5030, Zymo Research) according to the manufacturer’s protocol. This was followed by a second strand synthesis reaction and then Splinted Ligation Adapter Tagging (scSPLAT) with specially designed and pre-annealed oligos^85^. PCR was performed with TruSeq Unique Dual Indices (PN, 20040870, Illumina). Library quality control was performed on an Agilent 4200 TapeStation. Libraries were sequenced on an Illumina NovaSeq X instrument (150 bp PE reads).

Sequence data quality control, alignment, and generation of cytosine methylation reports: Binary base call (BCL) files from mTreg and xTreg sequencing libraries were converted to FASTQ format using bcl2fastq (Illumina). FASTQ files from mTreg libraries were adapter and quality trimmed using TrimGalore (v.0.6.10) with the following parameters specified: --non_directional --rrbs -- paired. FASTQ files from xTreg libraries were processed using TrimGalore (v.0.6.4). After adapter and quality trimming, 15 bases were further trimmed off at the 5’ end and 3’ end of the sequencing reads. FastQC (v.0.11.9) was used to assess the effect of trimming and overall quality distributions of the data.

Alignment to the *M. musculus* reference genome (mm10) was performed using Bismark (v0.22.3). We first prepared the *M. musculus* genome (mm10, downloaded from genome.UCSC.edu, last modified 2021-04-12) using *bismark_genome_preparation* with the -- bowtie2 option set. Next, we used *bismark* to map paired reads to the mm10 bisulfite genome with the non_directional option set. Next, we extracted the methylation information for every cytosine in the reads using *bismark_methylation_extractor* with the following parameters specified: -p -gzip -bedGraph. We then generated cytosine methylation reports using *coverage2cytosine* with the following parameters specified: --gzip --merge_CpG.

Identification of differentially methylated regions between mTregs and xTregs: We used DMRichR (v.1.7.1) to identify differentially methylated genomic regions between mTregs and xTregs. To this end, we first loaded the individual Bismark cytosine methylation reports for each sample into R. We then used the DMRichR::DM.R function (arguments: genome = “mm10”, testCovariate = “CellType”, minCpGs = 3, cutoff = 0.01) to identify differentially methylated regions. We considered regions with an adjusted *P*-value of < 0.05 as differentially methylated. We annotated genomic regions using the annotateRegions function from DMRichR with the TxDB parameter set to EnsDb.Mmusculus.v79 (version 2.99.0) and the annoDB parameter set to org.Mm.eg.db (version 3.20.0).

### Chromatin Immunoprecipitation and Quantitative PCR

Chromatin immunoprecipitation was performed by modifying a previously described protocol^86^. Briefly, splenic CD4 T cell subsets were enriched by MACS columns and sorted by FACS as described above, washed and resuspended in DMEM with 10% FBS medium as single cell suspensions. After fixing these cells in 1% formaldehyde in PBS for 15 min at RT on a horizontal rotator, the cells were sheared by sonication to generate chromatin fragments. Sample supernatant containing the DNA was quantified and 2% saved as input. PureProteome^TM^ Protein G Magnetic Beads (Millipore) were pre-incubated with 0.5 mg/ml anti-Foxp3 antibodies before incubating overnight with 3.4 µg of the chromatin fragments. After washing, the immune complexes were eluted from the beads, and the protein-DNA crosslinks reversed by incubating at 65°C overnight. After treatments with RNase, followed by proteinase K, the samples were purified with MicroChIP DiaPure columns (catalog no. C03040001, Diagenode) and eluted in 10 mM Tris-HCl elution buffer (pH 8.5) prewarmed to 55°C. Quantitative (q) PCR was used to evaluate the relative abundance of Foxp3 binding to the CNS2 region of the *Foxp3* gene in the input and FOXP3-ChIP DNAs. qPCR assays were performed in triplicate using Maxima^TM^ SYBR Green ROX mix (Fermentas) on thermocycler (Bio-Rad CFX96 C1000 Touch). The input DNA that lacked FOXP3 binding was used as control for normalization. The 2-ddt method was used for calculating enrichments for each target in ChIP-DNA relative to the input. The following previously published primers^87^ were employed: *Foxp3* promoter-F: 5’- CAATGCTGTCTCTACCTGCCTCG-3’; *Foxp3* promoter-R: 5’- CTCACCACAGAGGTAAAAGGTATCAATGA-3’; CNS2-F: 5’-GTTGCCGATGAAGCCCAAT-3’; CNS2-R: 5’-ATCTGGGCCCTGTTGTCACA-3’.

### Statistical analysis

All statistical tests were performed using GraphPad Prism (v6-v10) and presented in the corresponding figure legends.

## Data availability statement

All raw sequencing data generated in this study can be found in the Sequence Read Archive (SRA) at the NCBI National Library of Medicine (https://www.ncbi.nlm.nih.gov/sra) under the BioProject code: PRJNA1276081. Archived scripts and output files as at time of publication are available at Zenodo (doi:10.5281/zenodo.15632586).

## Acknowledgements

We would like to thank Drs. Alexander Rudensky (Memorial Sloan Kettering), John Harty (University of Iowa), and Rick Tarleton (University of Georgia) for friendly discussions, Dr. Alexander Rudensky for the FOXP3-DTR^GFP^ mice, Dr. William Heath (The Peter Doherty Institute for Infection and Immunity) for the PbT-II mice, Drs. Sungdae Park, Demba Sarr, and Douglas Menke (University of Georgia) for assistance with CHIP-qPCR assays, as well as the UGA CTEGD Flow Cytometry Core, UGA CVM Cytometry Core, UGA Biomedical Microscopy Core, and the UGA CTEGD animal research facility staff for technical support. These studies were funded by the US National Institutes of Health (AI168307, AI193123 to SPK, K99AI177948 to AAR). The Mali cohort study was funded by the Division of Intramural Research, National Institute of Allergy and Infectious Diseases, National Institutes of Health (PDC). The contributions of the NIH authors were made as part of their official duties as NIH federal employees, are in compliance with agency policy requirements, and are considered Works of the United States Government. However, the findings and conclusions presented in this paper are those of the authors and do not necessarily reflect the views of the NIH or the U.S. Department of Health and Human Services. SPK is a Burrows Wellcome Fund investigator.

## Author contributions

Conceptualization: NAEC-C, SPK; methodology: NAEC-C, AAR, NO-A, KDK, PDC, SPK; investigation: NAEC-C, AAR, CB, NO-A, DBRP, MRH, BT; writing: NAEC-C, AAR, SPK; funding acquisition: PDC, SPK; supervision: PDC, SPK.

---

**Figure S1.**
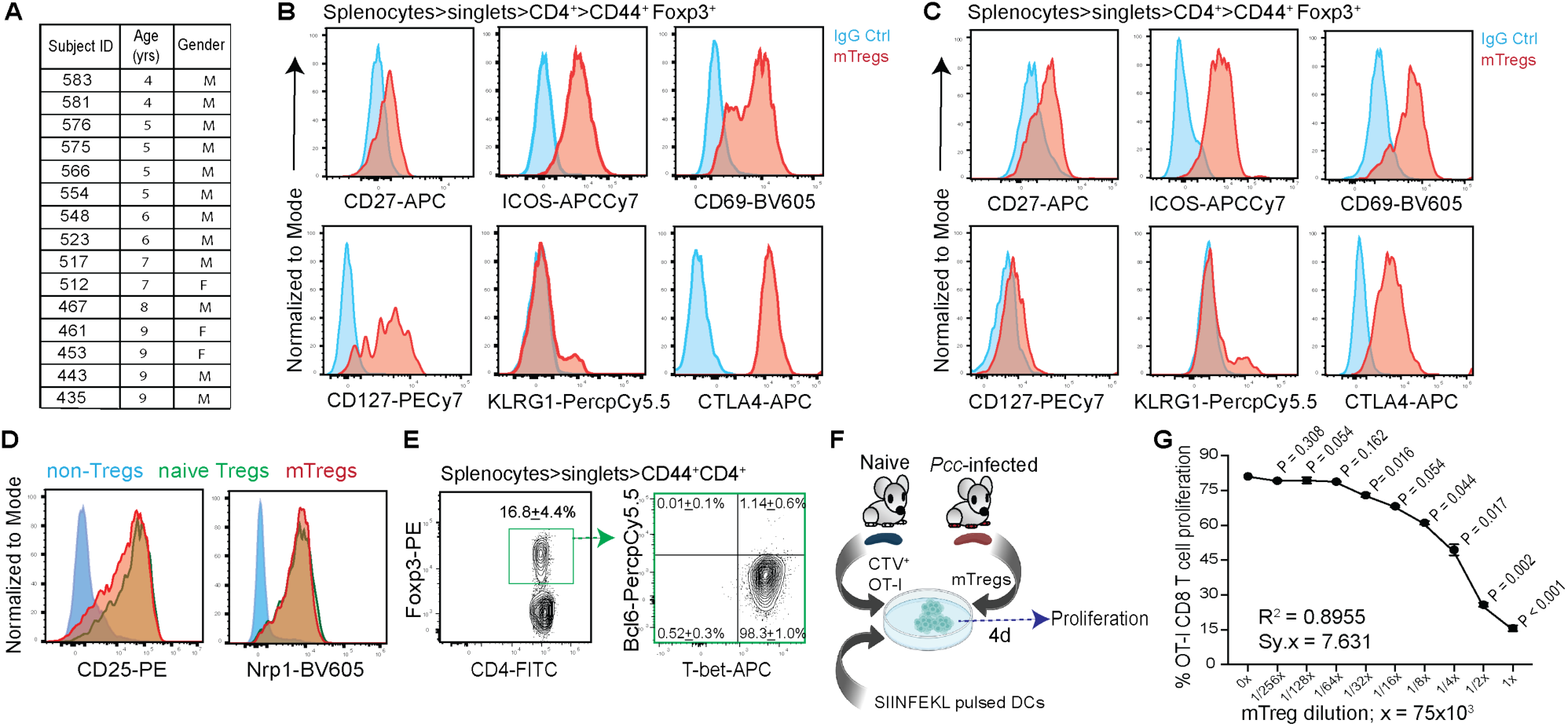
Phenotypic and functional characterization of memory Tregs. (A) Table depicting demographics of the Malian study subjects in Figures 1B, 2B. (B-C) Representative histograms (red) depicting expression of the indicated markers in splenic mTregs identified as in Figure 1D, but from *Listeria monocytogenes* (*Lm*)*-*infected (81d p.i, B) or Influenza A Virus (IAV, x31)-infected (42d p.i, C) B6 mice; blue histograms show IgG staining controls, from 1 of 3 experiments, n=3. (D) Representative histograms showing expressions of CD25 (IL-2Rα) or Neuropilin 1 (Nrp1) in splenic naïve Tregs (Foxp3^+^ CD4^+^) from naïve B6 mice, or the non-Tregs (Foxp3^-^ CD44^+^ CD4^+^) or mTregs obtained from *Pcc*-exposed B6 mice (181d p.i.). Representative data from 1 of 2 experiments, n=3. (E) Representative flow plots depicting T-bet or Bcl6 expression in mTregs obtained from *Pcc*- experienced B6 mice, 92d p.i. Numbers inset indicate gated cell frequencies presented as mean ± SD from 1 of 2 experiments, n=3. (F-G) B6 Dendritic cells (DCs) pulsed with chicken ovalbumin (OVA) antigen were co-cultured with CellTrace Violet (CTV)-stained TCR-transgenic OT-I CD8 T cells specific to OVA in the presence of increasing numbers of mTregs flow-sorted from *Pcc*-exposed (180d p.i.) FOXP3- DTR^GFP^ mice (F). Percentage of proliferated OT-I cells determined by CTV dilution; all cells isolated from the spleen, representative data from 1 of 3 experiments presented as mean ± SD from 3 technical replicates, analyzed using t-tests comparing each dilution to undiluted (0x) mTregs to obtain the indicated *P*-values (G).

**Figure S2.**
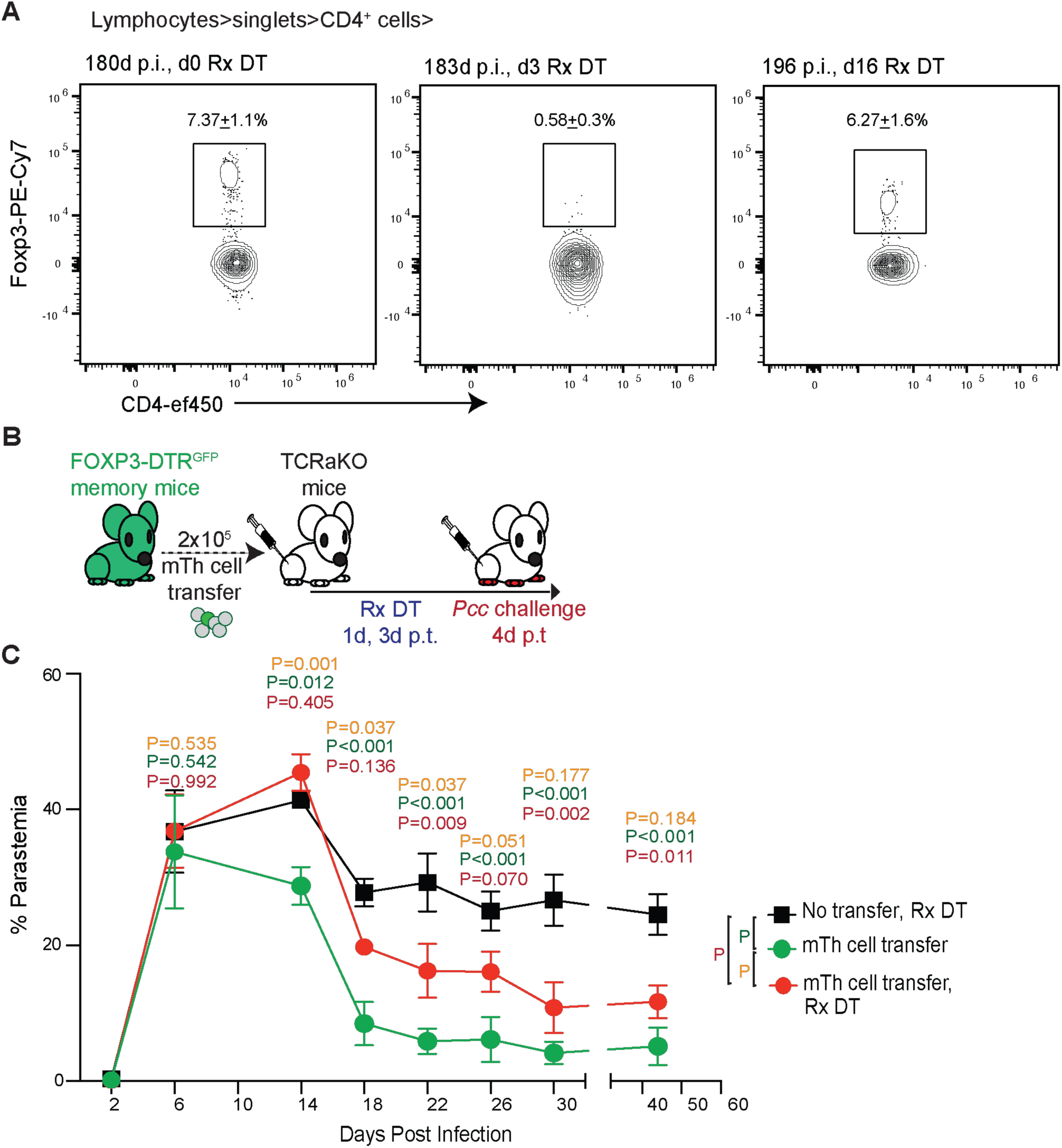
Memory Tregs essential for the transfer of immunity to malaria by CD4 T cells. (A) Representative flow plots depicting frequencies of Foxp3^+^ CD4 T cells in *Pcc*-experienced FOXP3-DTR^GFP^ mice at 0d p.t., 3d p.t. (183d p.i.) showing total Treg depletion and 16d p.t. (196d p.i.) showing Treg replenishment following DT treatments, as in Figure 2C. Numbers inset indicate the frequencies of the gated populations, presented as mean ± SD from 1 of 3 experiments; n=3. (B) 2x10^5^ memory CD4 T cells (mTh) from *Pcc*-experienced FOXP3-DTR^GFP^ mice (180d p.i.) were adoptively transferred to TCRaKO mice, which were then challenged with *Pcc* (5x10^5^ iRBCs/mouse) with or without DT treatment at the indicated timepoints (top panel). No cell transfer served as control. (C) Representative data on kinetics of parasitemia in the recipients from 1 of 3 experiments, presented as mean ± SD, analyzed using 2-way ANOVA corrected for multiple comparisons, yielding the indicated *P*-values color-coded for the groups; n=4 mice/group.

**Figure S3.**
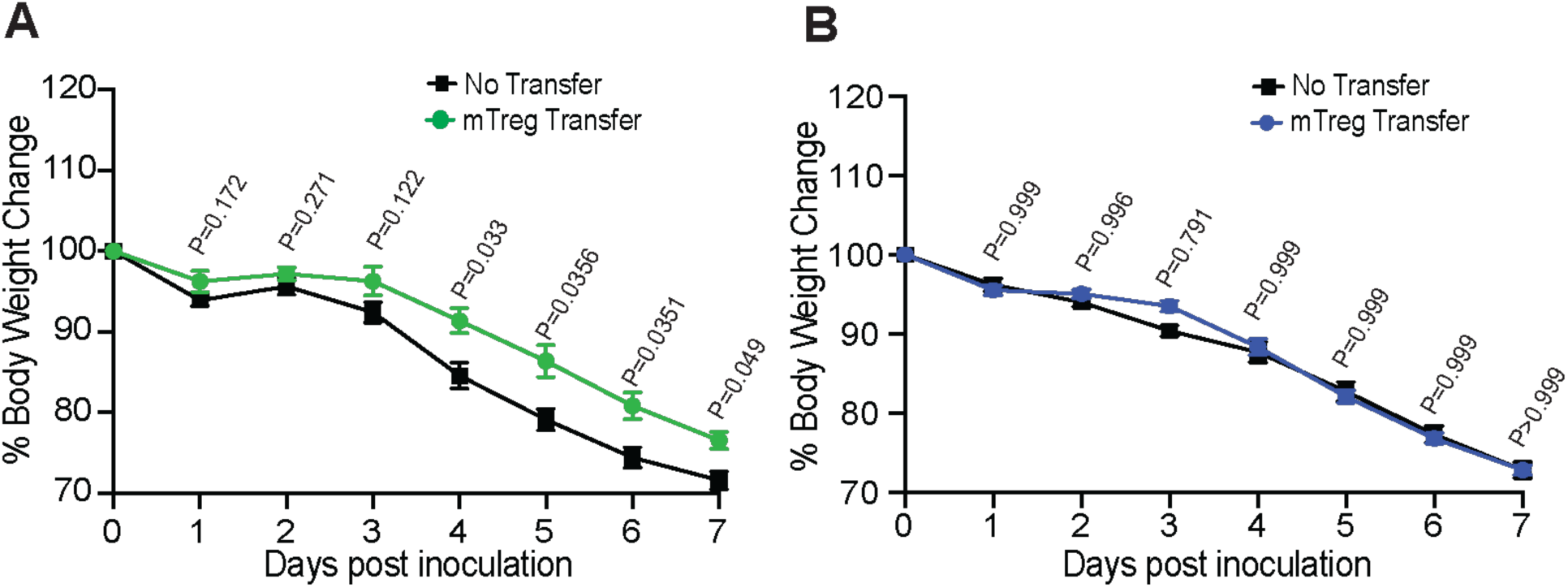
Memory Tregs from *Pcc*-exposed mice do not cross-protect from influenza. B6 mice receiving 2x10^5^ mTregs derived from IAV (x31, 49d p.i., A) or *Plasmodium* (*Pcc*, 180d p.i, B)-experienced FOXP3-DTR^GFP^ mice were challenged with IAV (500 PFU/mouse, PR8) at 1d p.t. Body weight change in the recipient mice until they reached humane end points determined. Combined data presented as mean ± s.e.m. from 2 independent experiments, analyzed using 2-way ANOVA with Sidak correction yielding the indicated *P*-values.

**Figure S4.**
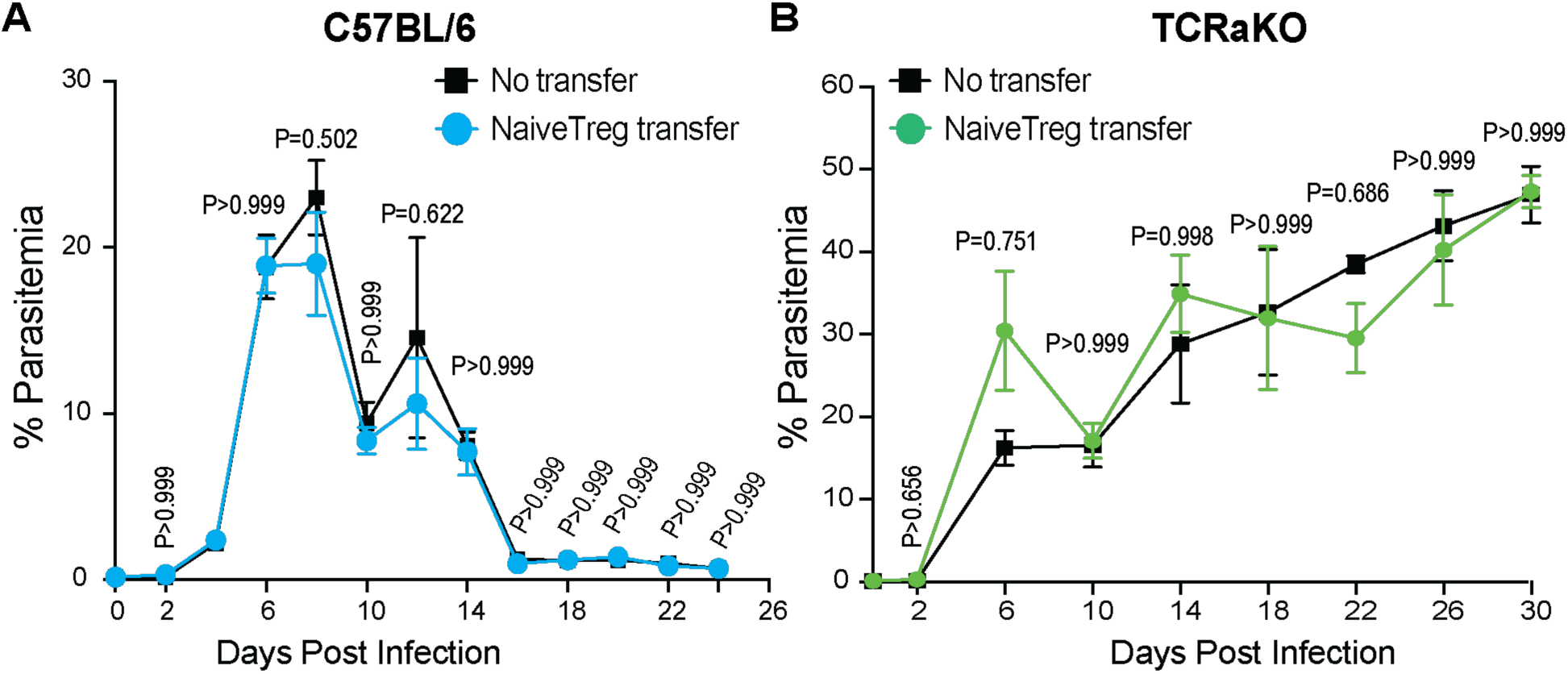
Naïve Tregs do not transfer protection against *Pcc* infection in mice. a-b, Kinetics of parasitemia in B6 (**A**) or TCRaKO (**B**) recipients of 2x10^5^ Tregs (Foxp3^+^CD4^+^) flow-sorted from naïve FOXP3-DTR^GFP^ mice. Data from 1 of 3 representative experiments (n=4), presented as mean ± SD, analyzed using 2-way ANOVA with Sidak correction yielding the indicated *P*-values.

**Figure S5.**
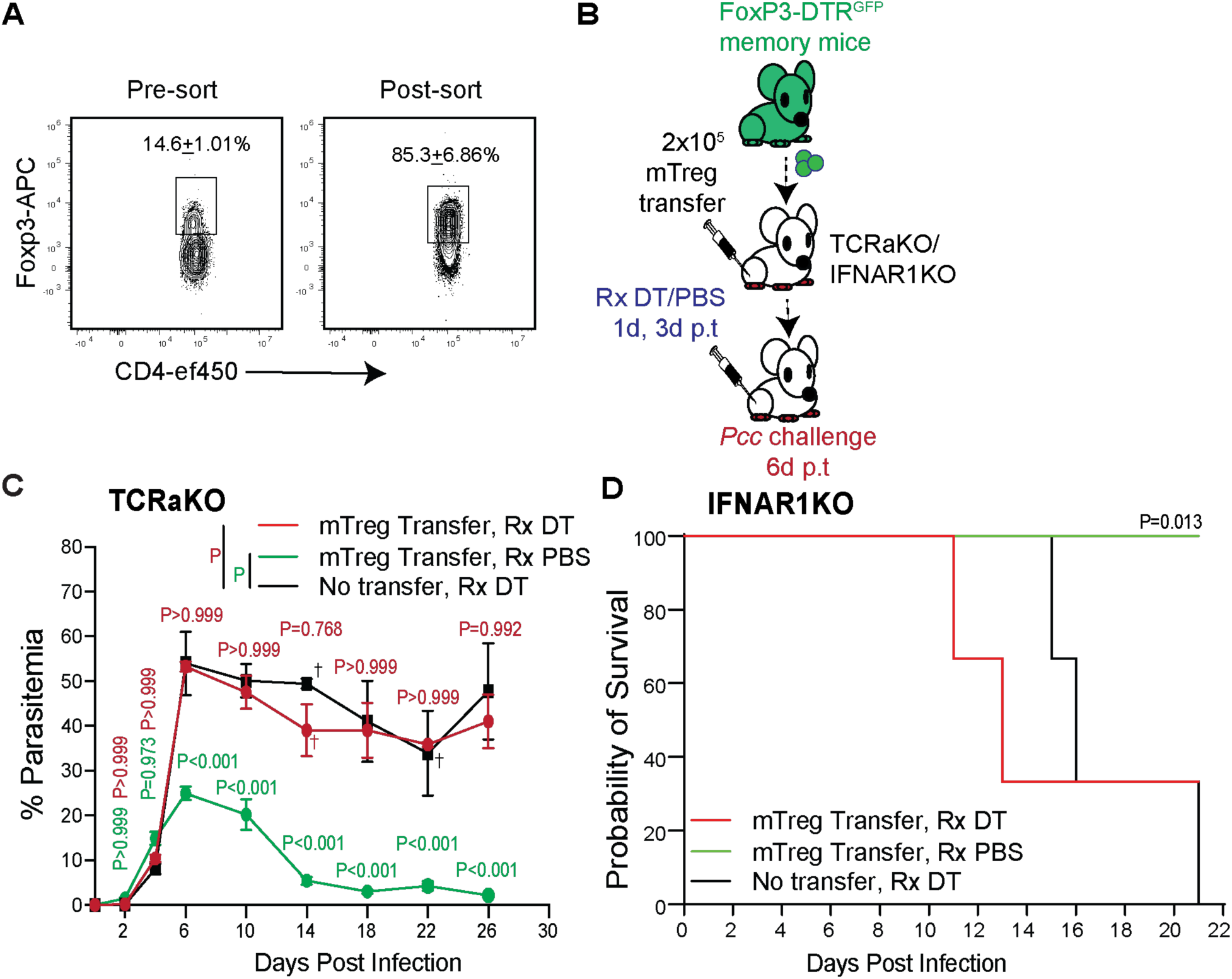
Depletion of memory Tregs before *Pcc* challenge reverses transfer of protection. (A) Representative flow plots showing the frequency of Foxp3^+^ CD44^+^ CD4 T cells in *Pcc*- experienced FOXP3-DTR^GFP^ mice (90d p.i.) before (left) or after (right) flow-sorting for their GFP expression. Numbers inset indicate the frequencies of the gated populations presented as mean ± SD from 1 of 3 separate sorting experiments, n=4 mice. (B-D) 2x10^5^ splenic mTregs from *Pcc*-experienced FOXP3-DTR^GFP^ mice (>180d p.i.) were adoptively transferred to TCRaKO or IFNAR1KO recipients, which were then treated with DT or PBS (1d, 3d p.t), and challenged with *Pcc* (5x10^5^ iRBCs/mouse) at 6d p.t. (B). Kinetics of parasitemia in TCRaKO (C) and survival in IFNAR1KO (D) recipients. Data presented as mean ± SD from 1 of 3 experiments, analyzed using 2-way ANOVA with Sidak correction (started with n=4 recipients, C) or Mantel-Cox test (started with n=3 recipients, D) yielding the indicated *P*-values. ^†^Death of 1 mouse/group.

**Figure S6.**
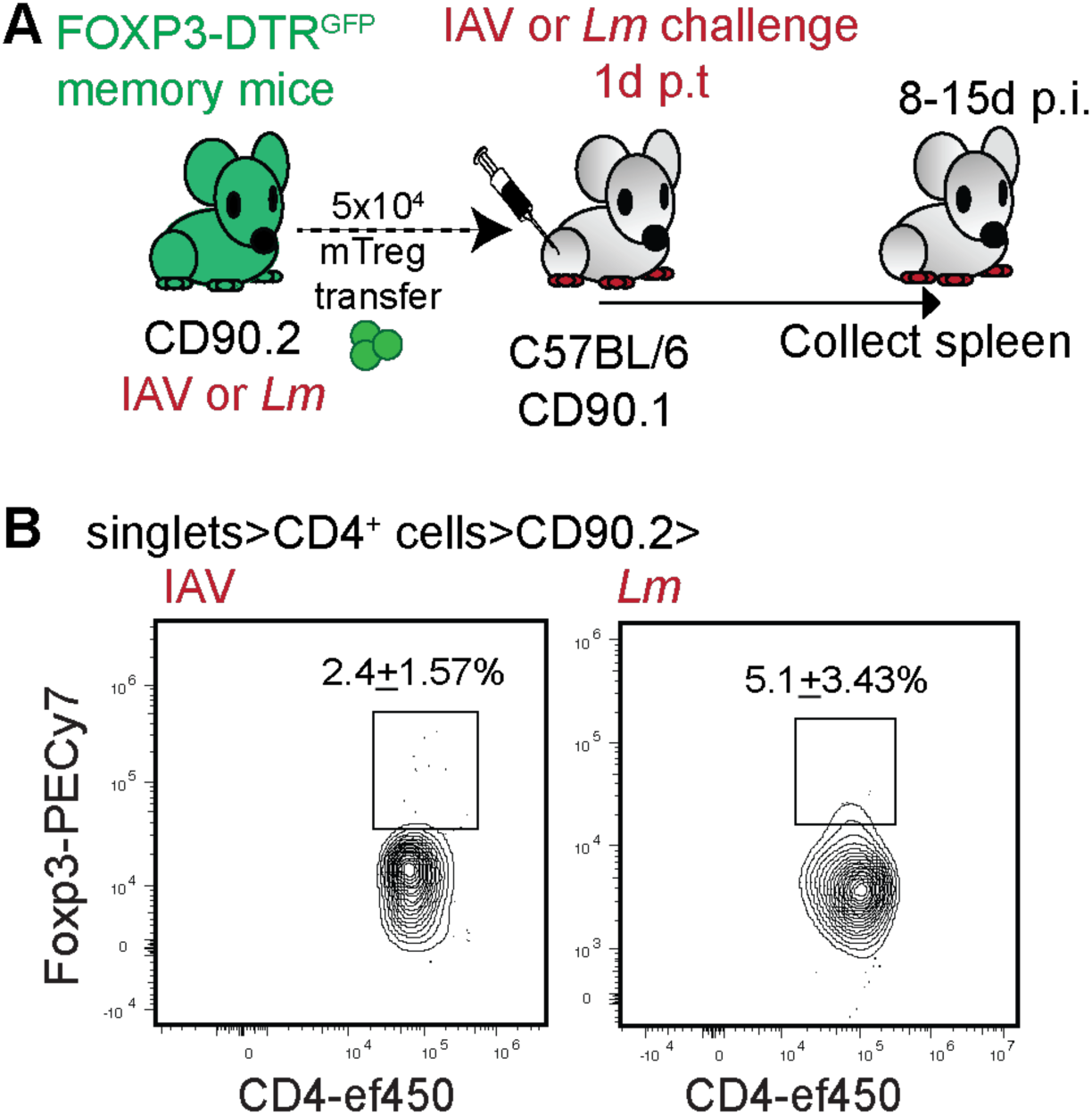
Memory Treg recall in *Lm* and IAV infections associated Foxp3 downregulation. (A) Experimental scheme: 5x10^4^ mTregs derived from IAV (x31, 49d p.i.) or *Lm* (81d p.i.)*-* experienced FOXP3-DTR^GFP^ mice (CD90.2) were adoptively transferred to congenically distinct (CD90.1) naïve B6 mice, the recipients challenged with IAV (PR8) or *Lm* and their spleens analyzed at 8d (IAV) or 15d (*Lm*) p.i. (B) Representative flow plots indicating Foxp3 expression in the IAV (left) or *Lm* (right)- experienced donor cells recovered from recipient spleens. Numbers inset show the gated cell frequencies presented as mean ± SD from 1 of 2 identical experiments, n=3 recipients.

**Figure S7.**
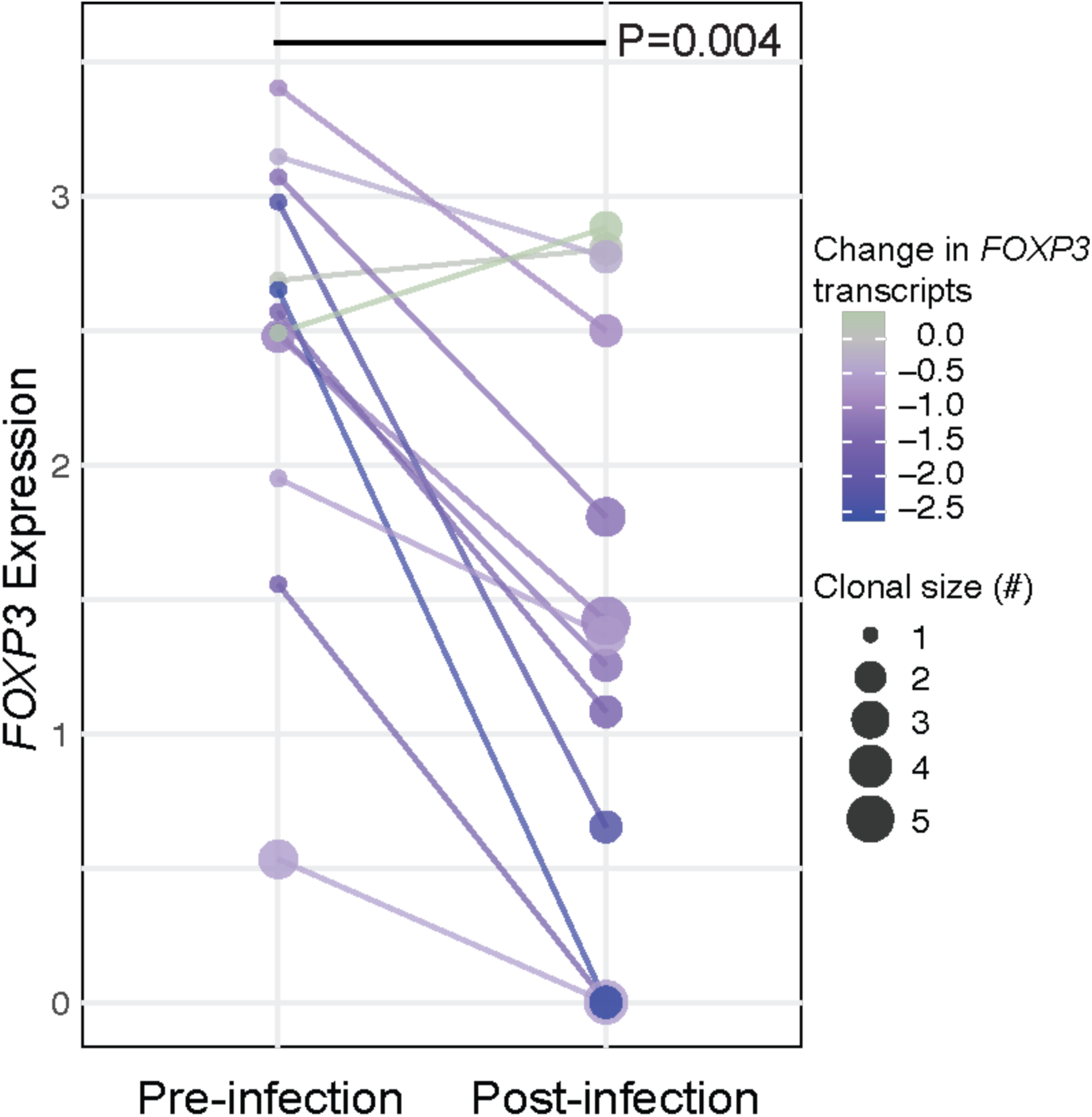
Recall of mTregs in humans is associated with FOXP3 downregulation. Combined data indicating *FOXP3* transcript levels in distinct Treg clones identified in human patient samples (n=4, PBMCs) longitudinally analyzed before or after (≥6d) *P. falciparum* infection. Individual clones indicated with the connecting lines. Primary scRNAseq data from Nideffer *et al*^21^, analyzed using paired Wilcoxon test to obtain the indicated *P-value*.

**Figure S8.**
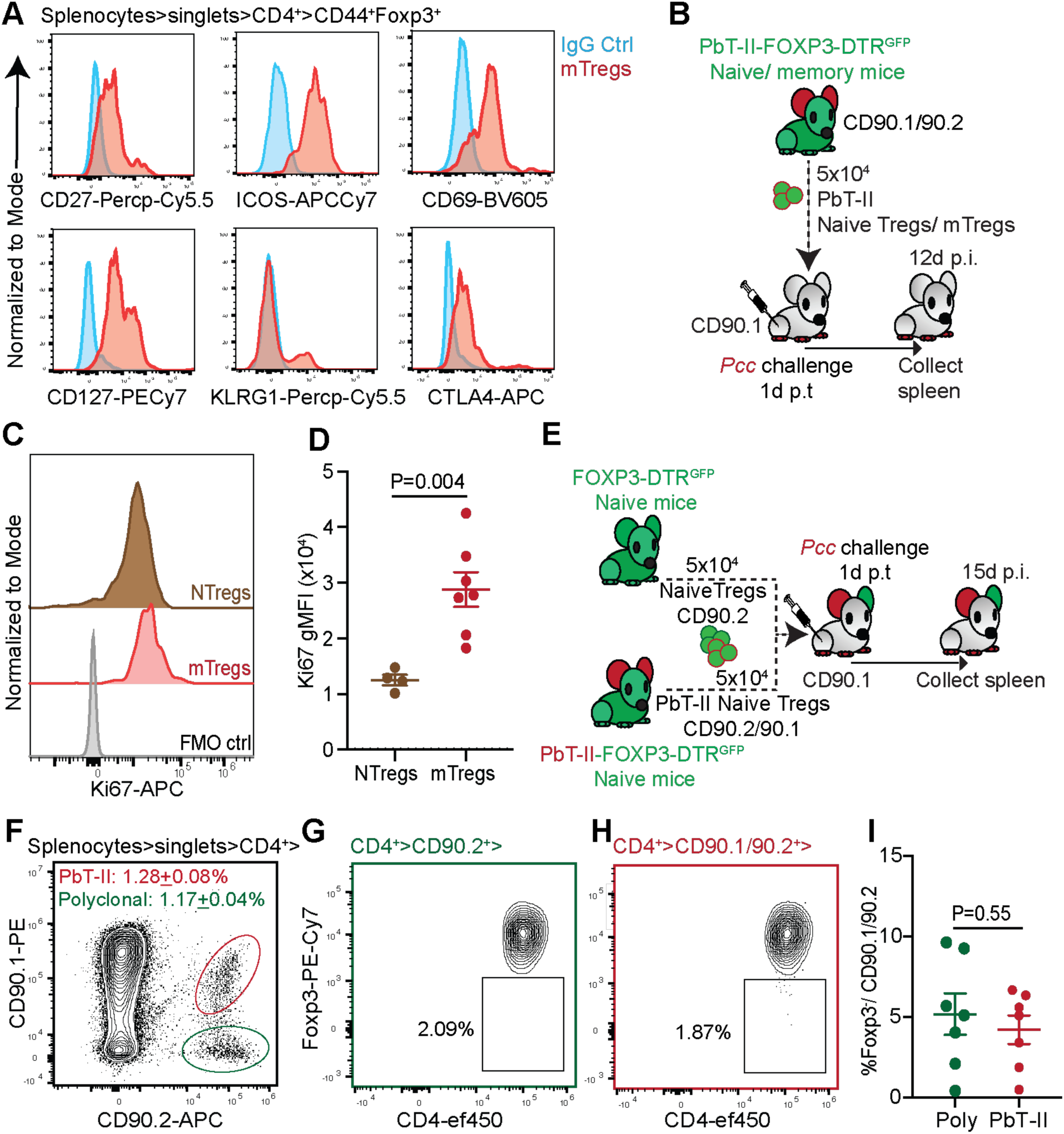
Naïve Tregs retain Foxp3 expression following antigen-induced proliferation. (A) Representative histograms showing key phenotypic markers of memory in the PbT-II mTregs generated as in Figure 4D. (B-D) 5x10^4^ PbT-II (CD90.1/90.2) naïve Tregs or mTregs were adoptively transferred to congenically distinct B6 mice (CD90.1) and challenged with *Pcc* iRBCs, analyzed at 12d p.i. (B). Staggered histograms indicating Ki67 expression in the donor cells isolated from recipient spleens (C); scatter plots summarize Ki67 expression (gMFI). Dotted line: pre-transfer mTregs, combined data from 2 separate experiments presented as mean ± s.e.m. analyzed using t-test to yield the indicated *P*-values with each dot representing a recipient (D). (E-I) 5x10^4^ polyclonal (CD90.2) and PbT-II-specific (CD90.1/90.2) naïve Tregs were adoptively co-transferred to congenically distinct (CD90.1) B6 mice, which were then challenged with *Pcc* (5x10^5^ iRBCs/mouse) and analyzed at 15d p.i.. Representative flow-plots show donor cells in recipient spleens (F) and Foxp3 expression in them (G, H) with the numbers inset indicating the frequencies of the gated population. Data summarized as mean ± s.e.m. from 2 independent experiments, analyzed using t-test to obtain the indicated *P*-value (I).

**Figure S9.**
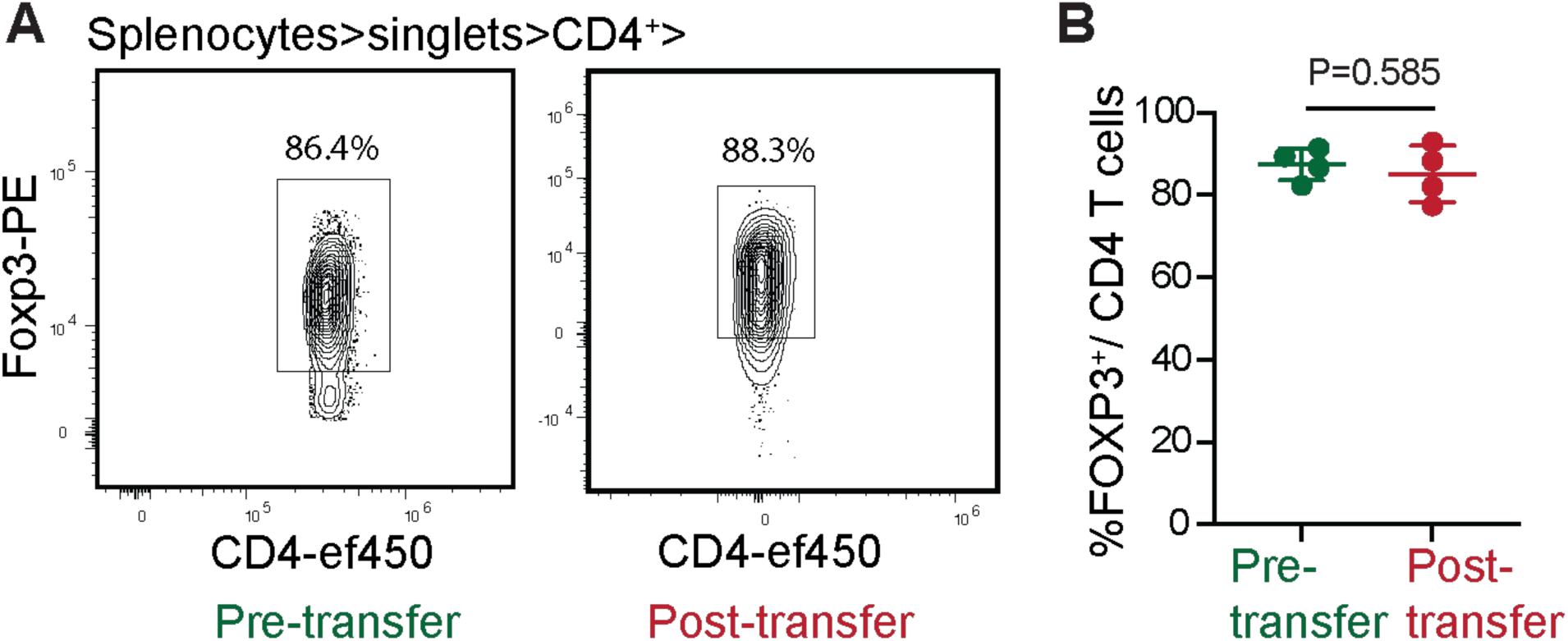
Memory Tregs retain Foxp3 expression during homeostatic proliferation. (A) Representative flow plots indicating the Foxp3-expressing cell subset within the sorted GFP^+^ PbT-II mTregs prior to adoptive transfer (*left*) to RAG2KO recipient mice and within the splenic CD4 T cell compartment of the recipients (*Right*), 11d p.t. (B) Summary data from 1 of 2 replicate experiments with each dot representing a mouse; n=4 recipients

**Figure S10.**
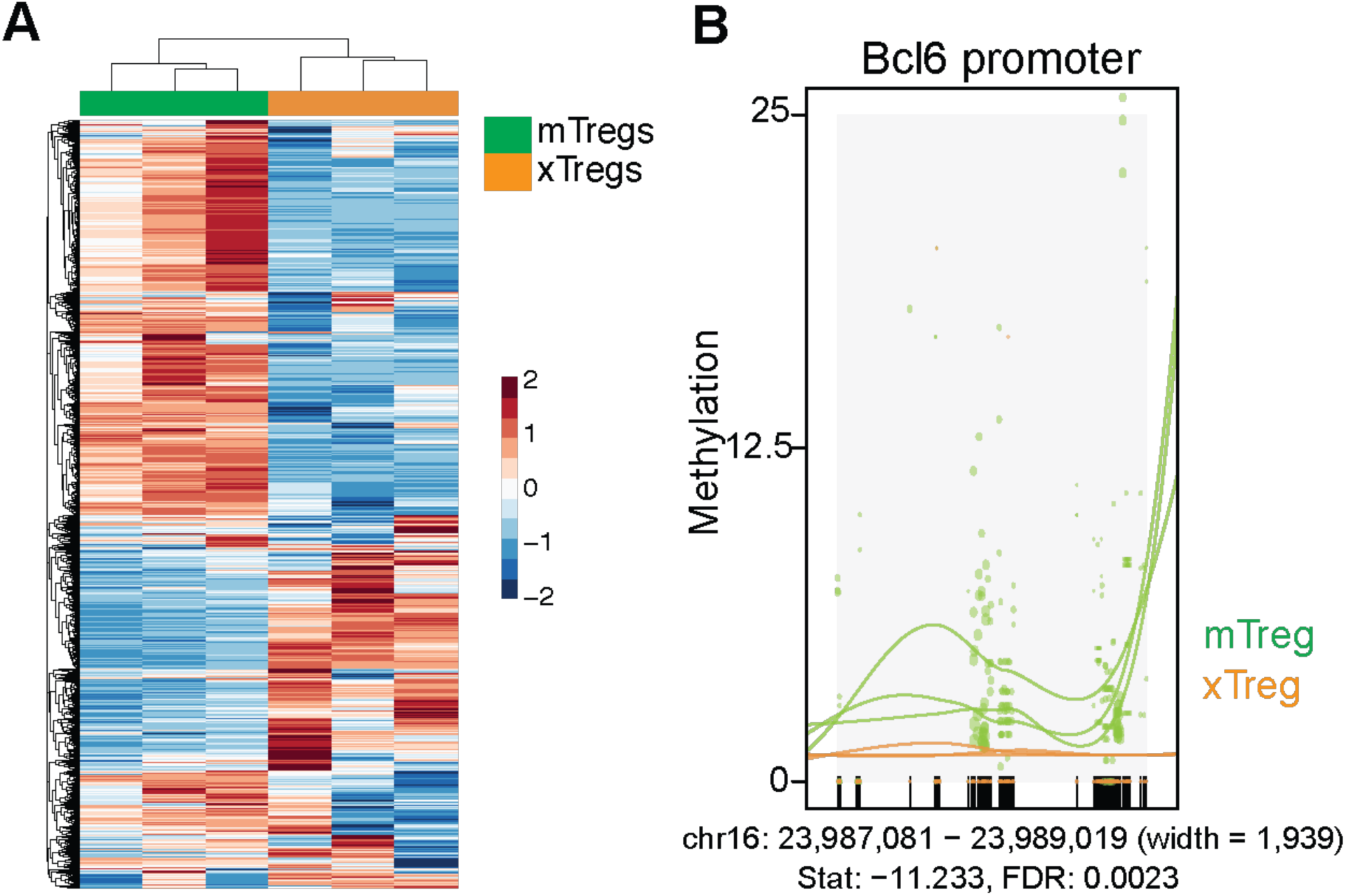
Differential methylation in mTreg and xTreg genomes. (A) Hierarchically clustered heatmap displaying differential DNA methylation patterns in mTregs and xTregs. Rows represent genome sites and columns represent WGBS samples (n = 3 for mTreg, n = 3 for xTreg). Color intensity reflects the relative level (scaled z-score) of methylation in each cell type. (B) Relative methylation levels across the *Bcl6* promoter in mTreg and xTreg genomes. Dot size corresponds to relative read depth, and each color-coded line connecting methylated loci represents an individual biological replicate (n=3 per cell type).

**Figure S11.**
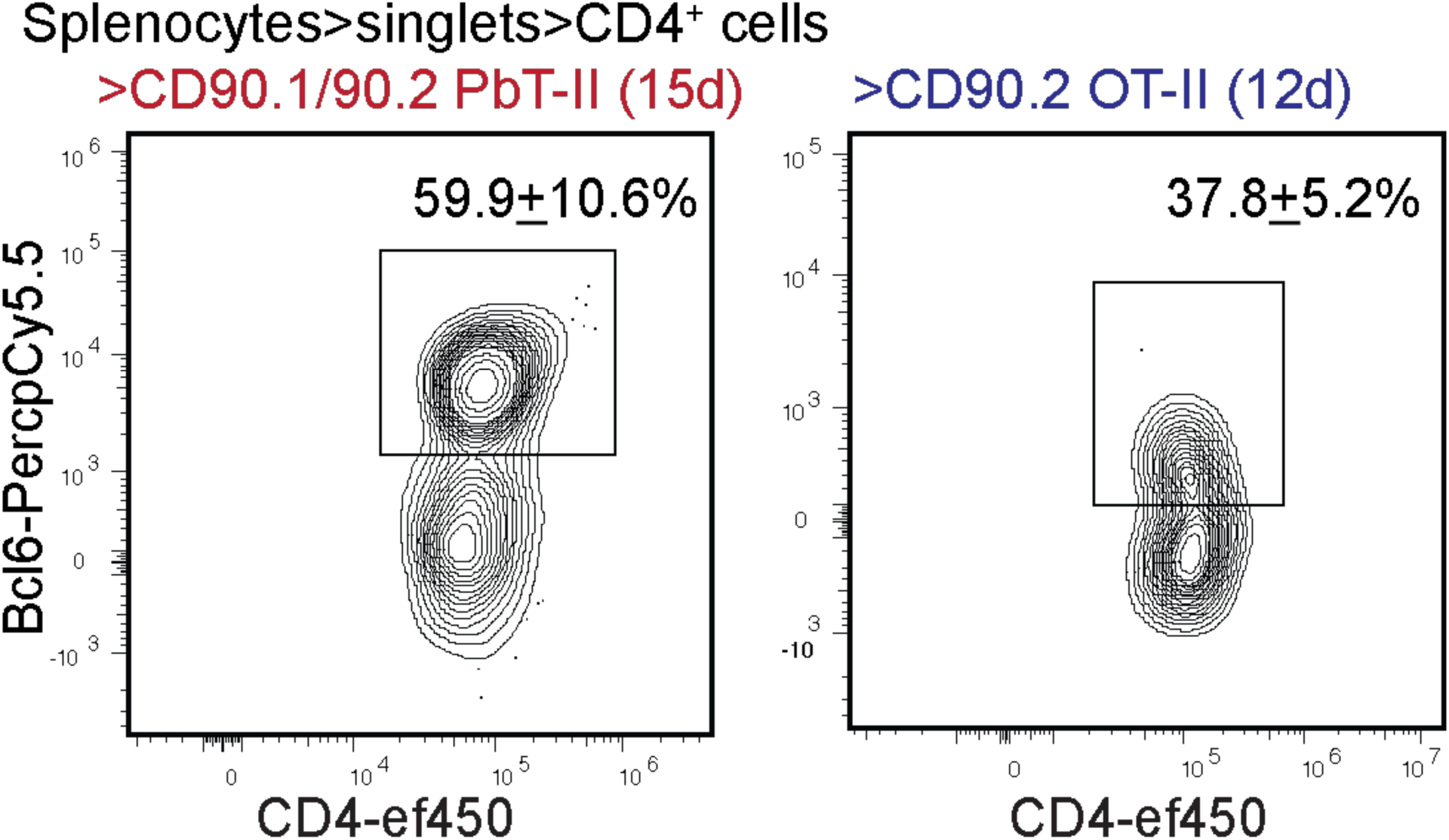
Bcl6 expression in PbT-II and OT-II mTregs. Representative flow plots showing Bcl6 expression in the adoptively transferred PbT-II (*Left*) or OT-II (*Right*) mTregs obtained from recipient spleens following *Py* or *Lm-Ova* challenge, as depicted in Figures 4D or 4H respectively. Numbers inset indicate the gated cell frequencies, combined data presented as mean ± s.e.m. from 2 replicate experiments; n=3 recipients/experiment.

**Figure S12.**
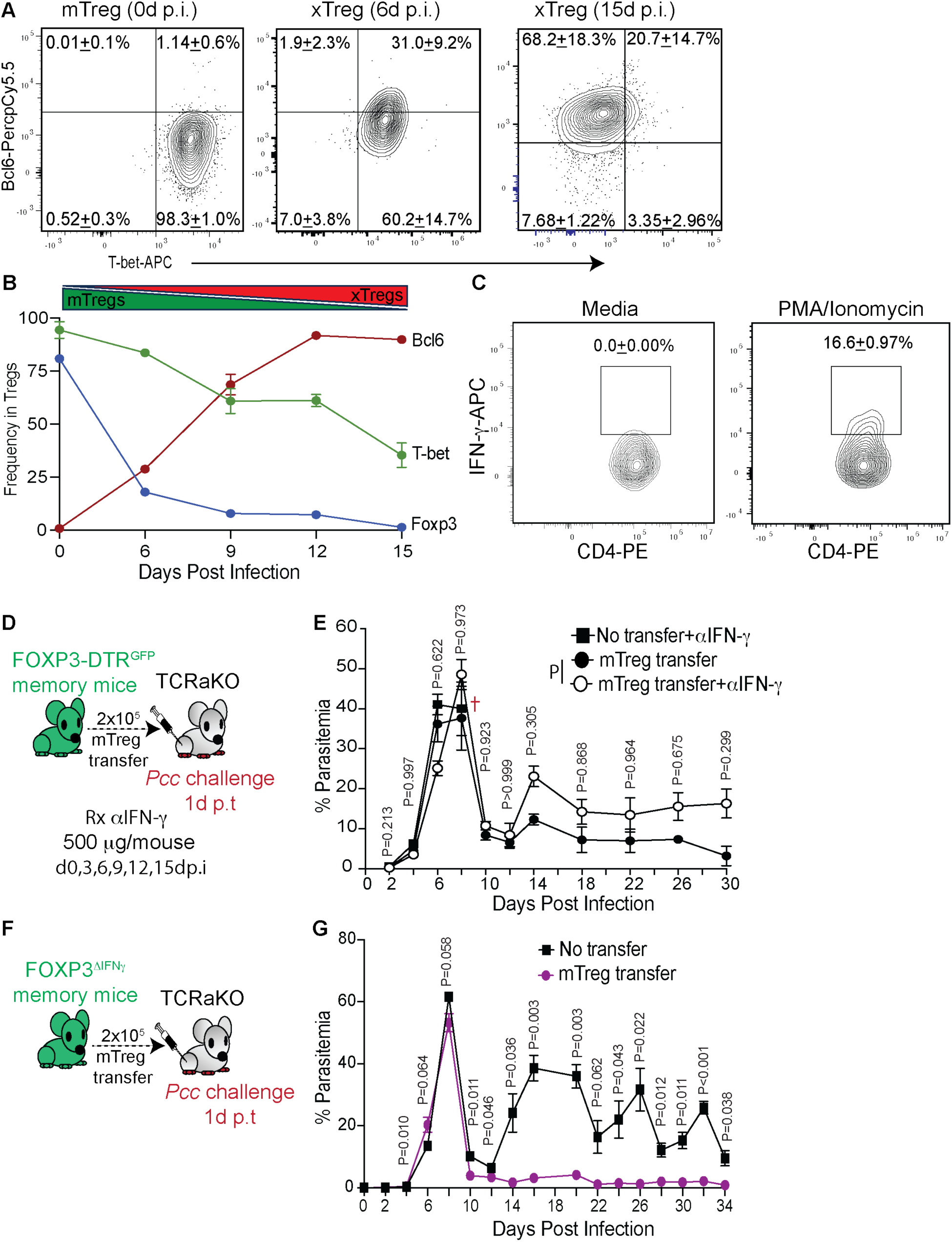
**IFN-**γ **produced by xTregs not essential for protection from *Pcc* challenge.** (A) Representative flow plots showing Bcl6 and T-bet expression in the xTregs obtained from the *Pcc*-infected mice generated as in Figure 5a, at the indicated days p.i. (B) Summary line graph depicting the kinetics of Foxp3, Bcl6 and T-bet expression in the adoptively transferred mTregs in the *Pcc*-infected recipient mice generated as in Figure 5a. Data presented as mean ± SD from 3 replicate experiments with n≥3 recipients/experiment. (C) Representative flow plots showing IFN-γ production in control (media) or PMA/ionomycin-stimulated xTregs obtained from *Pcc*-infected recipient mice, 16d p.i. (D-E) 2x10^5^ mTregs from *Pcc-*experienced FOXP3-DTR^GFP^ mice were adoptively transferred to TCRaKO mice, the recipients challenged with *Pcc* (5x10^5^ iRBC/mouse) and treated with control or anti-IFN-γ IgG at the indicated timepoints; TCRaKO mice which received no mTreg transfer served as control (D). Temporal kinetics of parasitemia in the recipients (E). (F-G) 2x10^5^ mTregs from *Pcc-*experienced FOXP3^βIFNγ^ mice were adoptively transferred to TCRaKO mice and the recipients challenged with *Pcc* (5x10^5^ iRBC/mouse) (F). Temporal kinetics of parasitemia in the recipient or control mice (G). (A, C) Numbers inset indicate the gated cell frequencies, presented as mean ± SD from 1 of 3 replicate experiments; n=3 receipt mice. (E, G) Data presented as mean ± SD from 1 of 3 replicate experiments, analyzed using t-tests at each timepoint to yield the indicated *P*-values. All cells were collected from the spleen for analysis or transfer.

**Figure S13.**
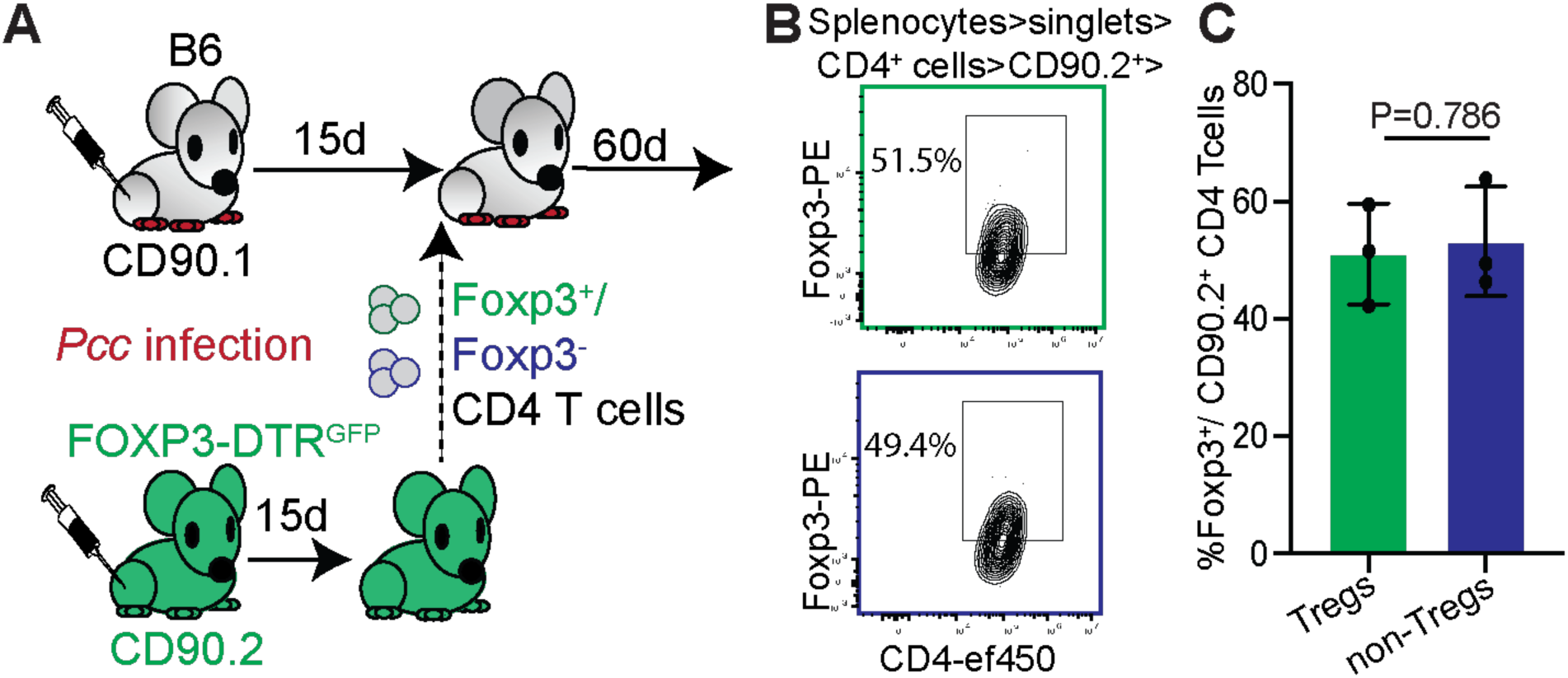
Memory Tregs can originate from Tregs and non-Tregs. (A) Flow-sorted splenic Foxp3^+^ (GFP^+^) or Foxp3^-^ (GFP^-^) CD4 T cells isolated from *Pcc-*infected FOXP3-DTR^GFP^ mice (CD90.2) at 15d p.i. were adoptively transferred to infection-matched (*Pcc*, 15d p.i.) congenically distinct B6 mice (CD90.1), which were then examined at 60d p.i. (B) Representative flow plots indicating Foxp3 expression within either donor CD4 T cell populations (Foxp3^+^: upper panel, Foxp3^-^: lower panel). c, Bar graph summarizing the data as mean ± SD from 1 of 3 replicate experiments with each data point representing a recipient, analyzed using t-test to yield the indicated *P*-value.

**Video S1. Adoptively transferred mTregs cluster with B cells in germinal centers**. Immunofluorescence microscopy and 3D volumetric reconstruction of the splenic architecture showing T cell zones (CD4^+^, pseudocolored blue), B cell follicles (B220^+^, pseudocolored red), GC reaction (GL7^+^, pseudocolored green) and the adaptively transferred mTregs (CD90.2^+^, pseudocolored grey) in *Pcc*-infected recipient B6 mouse at 15d p.i. The Tfh: B cell clusters containing the transferred cells within the GCs are clearly visible from 00:00:09s. Video representative of at least 3 regions observed in 5 sections examined from 3 separate mice from 3 separate experiments.

**Video S2. FOXP3-Cre^YFP^ mTregs generate robust Tfh:B cell clustering within the GCs.** Immunofluorescence microscopy and 3D volumetric reconstruction of the splenic architecture showing T cell zones (CD4^+^, pseudocolored blue), B cell follicles (B220^+^, pseudocolored red), GC reaction (GL7^+^, pseudocolored green) and the adaptively transferred FOXP3-Cre^YFP^ mTregs (CD90.2^+^, pseudocolored grey) in *Pcc*-infected recipient B6 mouse at 12d p.i. Tfh: B cell clusters within the GCs clearly visible from 00:00:10s. Video representative of at least 3 regions observed in 5 sections examined from 3 separate mice from 2 separate experiments.

**Video S3. FOXP3^ΔBcl6^ mTreg transfer fails to induce robust Tfh:B cell clustering within the GCs.** Immunofluorescence microscopy and 3D volumetric reconstruction of the splenic architecture showing T cell zones (CD4^+^, pseudocolored blue), B cell follicles (B220^+^, pseudocolored red), GC reaction (GL7^+^, pseudocolored green) and the adaptively transferred FOXP3^ΔBcl6^ mTregs (CD90.2^+^, pseudocolored grey) in *Pcc*-infected recipient B6 mouse at 12d p.i. Video representative of at least 3 regions observed in 5 sections examined from 3 separate mice from 2 separate experiments.

